# A common druggable signature of oncogenic CMYC, mutant KRAS and mutant p53 reveals functional redundancy and competition of the oncogenes in cancer

**DOI:** 10.1101/2023.12.20.572548

**Authors:** Maria Grześ, Akanksha Jaiswar, Marcin Grochowski, Weronika Wojtyś, Wojciech Kaźmierczak, Tomasz Olesiński, Małgorzata Lenarcik, Magdalena Nowak-Niezgoda, Małgorzata Kołos, Giulia Canarutto, Silvano Piazza, Jacek R. Wiśniewski, Dawid Walerych

## Abstract

Major driver oncogenes *CMYC*, mutant *KRA*S and mutant *TP53* often co-exist and cooperate in promoting human neoplasia. By CRISPR-Cas9-mediated downregulation we determined their proteomics and transcriptomics downstream programs in a panel of cell lines with activated either single or three oncogenes – in cancers of lung, colon and pancreas. This allowed to define and screen the oncogenes’ common functional program for anti-cancer target candidates, and find protocols which efficiently kill cancer cells and organoids by targeting pathways represented by a signature of three genes: *RUVBL1, HSPA9* and *XPO1*. We found that these genes were controlled by the driver oncoproteins in a redundant or competitive manner, rather than by cooperation. Each oncoprotein individually was able to upregulate the three target genes, while upon oncogene co-expression each target was controlled preferably by a specific oncoprotein which reduced the influence of the others. Mechanistically this redundancy was mediated by parallel routes of the target gene activation – as in the case of mutant KRAS signaling to C-JUN and GLI-2 transcription factors bypassing CMYC, and by competition – as in the case of mutant p53 and CMYC competing for biding to the target promoters. The transcriptomics data from the cell lines and patient samples indicate that the redundancy of the oncogenic programs is a broad phenomenon which may comprise even a majority of the genes dependent on the oncoprotein, as shown for mutant p53 in colon and lung cancer cell lines. Nevertheless, we demonstrate that the redundant oncogene programs harbor targets of efficient anti-cancer drug combinations, bypassing limitations of a direct oncoprotein inhibition.

## Introduction

The most deadly cancer types possess frequent upregulation of CMYC, mutations of *KRAS* and *TP53* ^1^. Yet, the knowledge about intersections of molecular programs driven by these common oncogenes is limited, and has not produced standard therapeutic solutions for cancer treatment ^2^.

The most frequently altered gene on average in human neoplasia is *TP53*, encoding an essential tumor suppressor p53, which upon acquisition of missense mutations not only loses its suppressive properties, but often acquires oncogenic gain-of-function ^3, 4^. Nevertheless, none of the drugs targeting mutant p53 directly is yet available as a standard therapy in patients ^5^, while the most advanced drug APR-246 is in phase 3 clinical trials ^6^.

KRAS missense mutations, occurring with high rates in pancreatic (more than 85% of cases), colorectal (∼40%) and NSCL (∼30%) cancers, lead to persistent activation of the protein’s GTP-ase activity and downstream signaling through PI3K-AKT-mTOR, MAP kinase and RAL pathways, among others ^7^. First and so far the only FDA-approved KRAS inhibitor for NSCLC treatment, sotorasib, targets the G12C mutation, however, according to the CodeBreaK trials this inhibitor does not contribute to improvement in patients overall survival ^8^. More inhibitors targeting other KRAS mutants are currently in trials ^8^.

One of the most often upregulated pro-oncogenic factors in cancer is *CMYC*, which is the master regulator of a vast transcriptional program crucial for cancer progression ^9^. *CMYC* expression alterations found in various types of cancers make this oncoprotein an appealing therapeutic target, but despite various strategies to inhibit *CMYC*, no clinical therapy is currently available ^10^.

While the described oncogenes are known to have independent activities, they represent important components of multistep carcinogenesis, where they functionally interact ^2^. KRAS is known to stabilize CMYC by activating the ERK1/2 which phosphorylates CMYC at serine 62 ^11^. Several studies demonstrated that mutant p53 and MYC may positively influence each other’s levels and activity ^12, 13^. Furthermore, transcriptional signature of mutant p53 in head and neck squamous cell carcinoma was enriched in MYC targets, and mutant p53 augmented CMYC binding to its target promoters ^14^. Analysis of mutant p53 and KRAS interactions was done in pancreatic ductual carcinoma (PDAC) often harbouring co-occurring *KRAS* and *TP53* mutations. Kim et al. ^15^ showed that KRAS-RAF-MEK-MAPK pathway activates CREB1, which is bound by mutant p53 to upregulate FOXA1 transcription factor program and PDAC metastasis. Another study revealed that mutant p53 rewires the splicing of GTP-ase-activating proteins to promote activation in KRAS signaling ^16^. Thus, the interactions described are limited to several mechanisms of cooperation between the trio of the oncogenes.

In this study, to increase knowledge on mutant p53, mutant KRAS and CMYC interactions and their downstream programs we systematically surveyed their transcriptomics and proteomics programs in a panel of colon, lung and pancreatic cancer cell lines. We defined common and specific pathways driven by the trio of oncogenes and found that simultaneous targeting of the common pathways can be efficient in killing cancer cells of various tissue origins, comprising a “druggable driver oncogene signature”. Interestingly, we discovered that these targets are not upregulated as a result of an oncogene cooperation, but rather their redundancy and competition – each driven dominantly by the oncogene which can be replaced, rather than augmented, by the presence of others. This implies that critical molecular pathways controlled by the major oncogenes possess a “safety mechanism” which guarantees activation in a background of any of the major oncogenic drivers. The robustness of oncogene redundancy described here makes these pathways a valuable target in attempts to kill cancer cells.

## Results

### Proteomics and transcriptomics reveal common, targetable pathways driven by mutant p53, mutant KRAS and MYC

To reveal protein and RNA populations controlled by oncogenes – mutant *TP53*, mutant *KRAS* and *CMYC* - we used CRISPR-Cas9-mediated editing to downregulate their expression levels in a panel of cell lines derived from cancer types frequently driven by the mentioned oncogenes (Fig. 1A). We stably introduced Cas9 to cell lines of lung and colon cancer which either harbor co-expressed missense hotspot mutant *TP53*, *KRAS* and oncogenic *CMYC* or each of the oncogenes individually. Additionally we performed the same procedure in a pancreatic cancer cell line PANC1, which, as in the majority of pancreatic ductal adenocarcinomas, contains coexpressed *KRAS* and *TP53* mutants, and hyperactive *CMYC* (Fig. 1A). In all cases pairs of gRNAs have been introduced by transient transfection to cell lines with the stable Cas9 to perform NHEJ-mediated knockouts of the oncogenes. Samples were collected at 48h post transfection and controlled for significant decrease of the oncogenes’ RNA and protein (Supplementary Figure 1A). Upon performing a label-free proteomics with >10K indentified protein sensitivity per sample, and a total RNA-seq (>60M reads), we carried out differential analyses between three biological replicates of non-specific gRNA control and gRNAs targeting each oncogene (Supplementary Table 1-2, Supplementary Fig. 1b-c). The results of this differential analysis, when included in the hierarchical clustering (5569 proteins and 15453 mRNA with quantified differences across all the samples), revealed that the change values cluster preferably with each of the oncogenes, rather than the tissue of origin or the cell line (Fig. 1 B, C). Hence, we proceeded to evaluate the downstream functionality of these molecular programs in the oncogene-focused manner.

**Figure 1.**
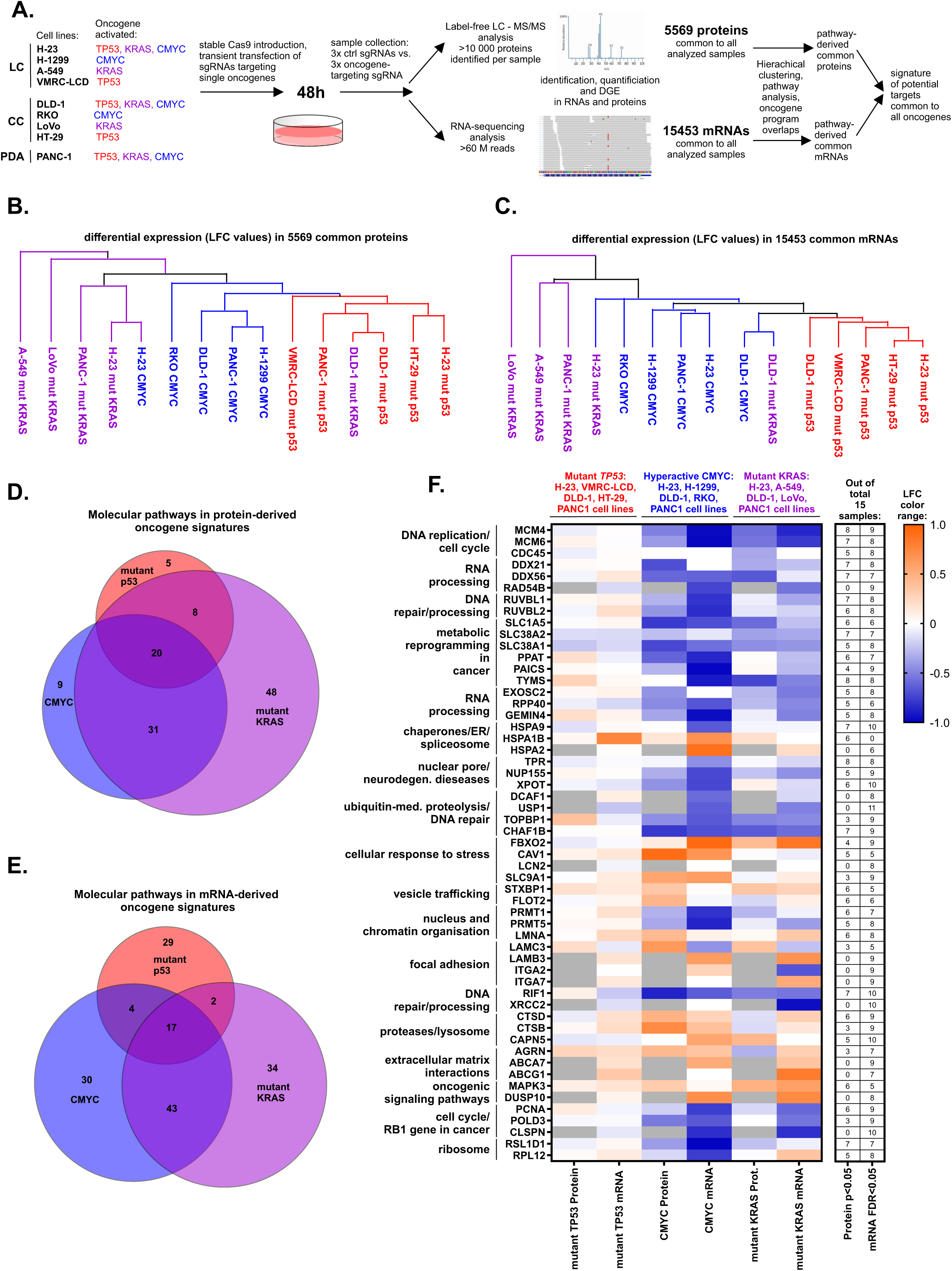
**A.** Experimental flow of the CRISPR-mediated oncogene editing in the indicated cancer cell lines (LC – lung cancer, CC – colon cancer, PDA – pancreatic ductal adenocarcinoma), followed by differential proteomics, transcriptomics and their subsequent analyses. **B.** Hierarchical clustering (Euclidean distance) of the differentially regulated 5569 proteins common to all the analyzed samples. Colors indicate programs dependent on each of the listed oncogenes. **C.** Hierarchical clustering performed as B, for the common 15453 mRNAs differentially regulated in all the samples. **D.** Venn diagram showing overlap of pathways significantly (FDR<0.05) associated by the Clue-GO software with the proteins differentially regulated by each of the three oncogenes indicated by names and colors across all the cell lines with a given oncogenes. The cutoff of p<0.05 for the differential analysis was used for the proteins included in the pathway association, followed by a duplicate filtration in each of the oncogenes’ programs. Significantly (p<0.05) up- and downdregulated proteins both were used in the pathway association. **E.** Overlap as in D. for mRNA-derived pathways regulated by the indicated oncogenes. **F.** A heatmap indicating average protein and mRNA level changes (Log Fold Change – LFC – range indicated at the color scale) of the listed genes derived from the common pathways in D and E, in cell lines with mutant *TP53*, mutant *KRAS* or hyperactive *CMYC*. The table on the right side shows how many times each of the listed genes was significantly changing level on the mRNA (FDR<0.05) and protein (p<0.05) levels in all the analyzed samples. The genes are assigned to pathway-derived functional groups shown on the left.

We fused the lists of protein and mRNA differentially regulated by each oncogene in all the cell lines it is present in, with a cutoff of p<0.05 for proteins and FDR<0.05 for mRNAs, filtered it for duplicates, and performed pathway analysis on each oncogene signature using Clue-GO software. The molecular pathways were then overlapped for proteins and mRNAs separately to understand specific and common aspects of the oncogene-driven programs (Fig. 1 D. E, Supplementary table 3). The analysis revealed a significant amount of common pathways, we further focused on the common programs of the oncogenes to hunt for potential universal downstream targets in cancer – both processes and individual proteins/genes. To extract these potential targets from the common pathways we scored how many times each protein or gene from the common pathways is present as significantly (p/FDR<0.05) changing level in each cell line. The protein/mRNA level change heatmap for the most frequently regulated proteins and mRNAs driven by the trio of the oncogenes is shown in Figure 1F. To be included in the heatmap the given gene had to be present in the common mRNA or protein pathways (Fig. 1D, E), and significantly change level in at least 6 of protein and mRNA readouts of the total 30 protein and mRNA analyses. As visible in the rightmost table in Fig. 1F none of the individual genes was significantly regulated in more than 17 of 30 total samples (15 for mRNA and 15 for proteins). The heatmap shows average changes of mRNA/protein levels (intensity) and change direction (color) in cell lines sharing each oncogene – demonstrating that mutant *TP53* is on average the least strong regulator of the target gene protein and mRNA change, compared to mutant *KRAS* and hyperactive *CMYC*. The genes have been assigned to functional groups (Fig. 1F) derived from the common oncogene driven processes (Supplementary table 3). As described in the next section these groups were targeted in an siRNA mini-screen to find specific vulnerabilities in the common molecular programs driven by mutant *TP53*, mutant *KRAS* and hyperactive *CMYC*.

### Targeting the signature controlled by mutant p53, mutant KRAS and overexpressed CMYC is efficient in killing lung, colon and pancreatic cancer cells

Altogether nineteen pathways, indicated in bioinformatics analysis as common to molecular programs controlled by mutant *TP53*, mutant *KRAS* and overexpressed *CMYC*, were targeted with sets of two to four siRNAs (Fig. 2A, Supplementary Table 4). siRNA mini-screen was performed in a panel of nine cell lines of colon (DLD1, LoVo, RKO), lung (H23, A549, VMRC-LCD) and pancreatic (PANC1, MIAPaCa2, BxPC3) cancers, as well as in two untransformed, normal human fibroblasts as control cells (F02, F03), with the aim to find functional gene groups whose depletion causes decrease of viability cancer cell lines and not of normal cells.

**Figure 2.**
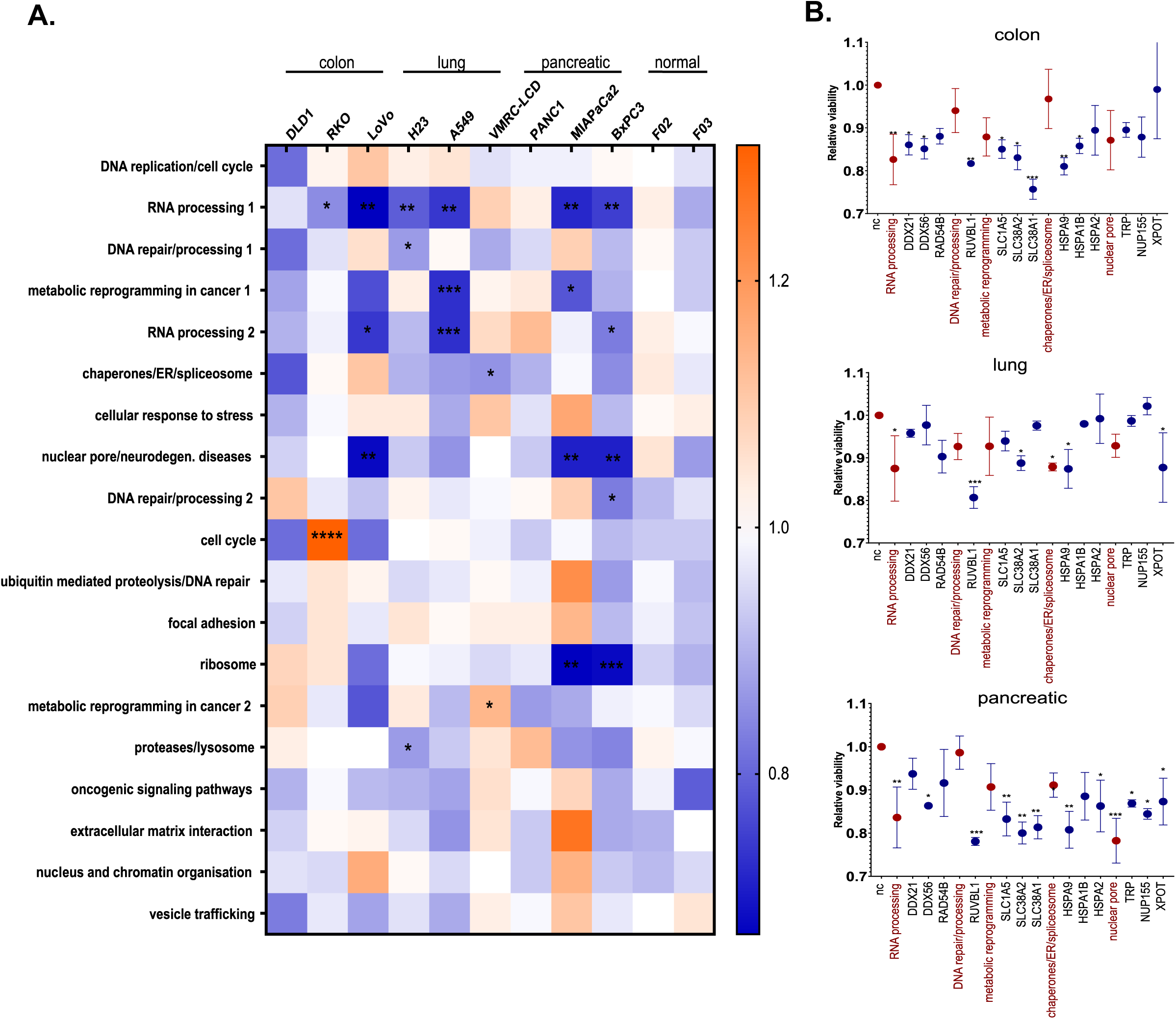
**A.** Colon, lung and pancreatic cancer cell lines and two normal fibroblasts lines were transfected with a mixture or 2-4 siRNAs, targeting genes belonging to functional groups (siRNAs listed in Supplementary Table 4). Resazurin assay was used for viability measurement 48h post transfection. Data in the heatmap are presented as means of n=2 biological replicates for each cell line and analyzed with One-way ANOVA (uncorrected Fisher’s LSD) versus siRNA negative control. **B.** Viability of colon (DLD1, RKO, LoVo), lung (H23, A549, VMRC-LCD) and pancreatic (PANC1, MIAPaCa2, BxPC3) cancer cell lines treated with single (dark blue) or combination of siRNAs (red) best performing in the siRNA mini-screen (A), targeting: helicase activity, ATPase activity, amino acids transport, chaperones and nuclear pore complex/transport. Viability was measured as in (A). Each result is a mean of n=6 (two biological replicates for each cell line), error bars represent SEM. Data were analyzed with One-way ANOVA (uncorrected Fisher’s LSD). A and B: *p < 0,05, **p<0,01, ***p<0,001.

Targeting helicases (DDX21, DDX56, RASD54B), amino acids transporters (SLC1A5, SLC38A2, SLC38A1) and nuclear pore complex/transport components (TPR, NUP155, XPOT) decreased cancer cell lines viability most significantly compared to normal cells, while we also observed the tendency to decrease viability across the tested cell lines in the case of blocking DNA repair/processing activity (RUVBL1, RUVBL2) and targeting chaperones (HSPA9, HSPA1B, HSPA2). Subsequently we tested the siRNAs from selected groups individually in order to determine depletion of which protein had the most significant impact on cancer cells survival (Fig. 2B, Supplementary Table 4). Obtained results suggested that siRNAs targeting *RUVBL1*, *HSPA9* and *SLC38A2* individually and nuclear pore transport as a group resulted in the strongest decrease of cancer cells viability and thus we concluded, that proteins encoded by these genes could lie at the intersection of vital programs of cancers driven by mutant *TP53*, mutant *KRAS* and hyperactive *CMYC* and as such be tested as therapeutic targets. Moreover, silencing of the oncogenes directly resulted in less significant decrease in viability of the same cancer cell line panel, especially in the case of the cell lines with the co-presence of the oncogenes (Supplementary Fig. 2A).

Subsequently we targeted proteins encoded by genes revealed in siRNA mini-screen as potential vulnerabilities common to mutant *TP53*, mutant *KRAS* and hyperactive *CMYC* programs. We tested the effect of CB6644 (targeting RUVBL1/2 complex), MKT077 (inhibiting HSPA9), selinexor (blocking XPO1, nuclear exportin – common inhibitor of nuclear transport mediated by TPR, NUP155 and XPOT) and MeAIB (targeting SLC38A2) on cell lines representing three different cancer types versus normal fibroblasts. After treating cell lines with increasing concentrations of inhibitors (Supplemetary Fig. 2B) we calculated for each cell line the half maximal concentration of inhibitor (IC_50_, Supplementary Table 5). Only for MeAIB we were not able to obtain IC_50_ form curves fitted with r^2^>0.9, as it did not affect significantly the viability of cancer cell lines, thus we decided to exclude this inhibitor from further experiments (data not shown). Each cancer cell line was treated with selected inhibitors used as single agents or in combination, with concentrations corresponding to the obtained IC_50_ values. For normal fibroblasts we used the concentration of each inhibitor representing the highest IC_50_ among all cancer cell lines.

Simultaneous XPO1 and HSPA9 targeting revealed high efficiency of this combination in killing cancer cells with insignificant impact on viability of the normal fibroblasts. The effect of significant decrease in viability after using combination of selinexor and MKT007 in comparison when only single inhibitor was used, was observed across all tested cancer cell lines (Fig. 3A). In the case of simultaneous blocking of RUVBL1/2 ATPase activity and inhibiting XPO1 results also showed high vulnerability of cancer cells to this combination. Ten out of twelve cancer cell lines presented significant viability decrease in comparison with the use of single agents (Fig. 3B). Importantly, treatment with each inhibitor as a single agent or as a mixture showed low toxicity to normal fibroblasts (Fig. 3C). We also tested the combination of MKT077 and CB6644, however it did not show statistically significant improvement in comparison to effect of single inhibitors in most of the tested cell lines (Supplementary Fig. 3A). Similarly, mixture of MKT077, CB6644 and selinexor did not turn out to be more efficient that pairs of inhibitors (Supplementary Fig. 3B). Nevertheless, the efferent double drug combinations (Fig. 3 A-B) had matching or significantly better performance in killing cancer cell lines than typical chemotherapy protocols used in colon, lung or pancreatic cancers (Supplementary Fig. 3C).

**Figure 3.**
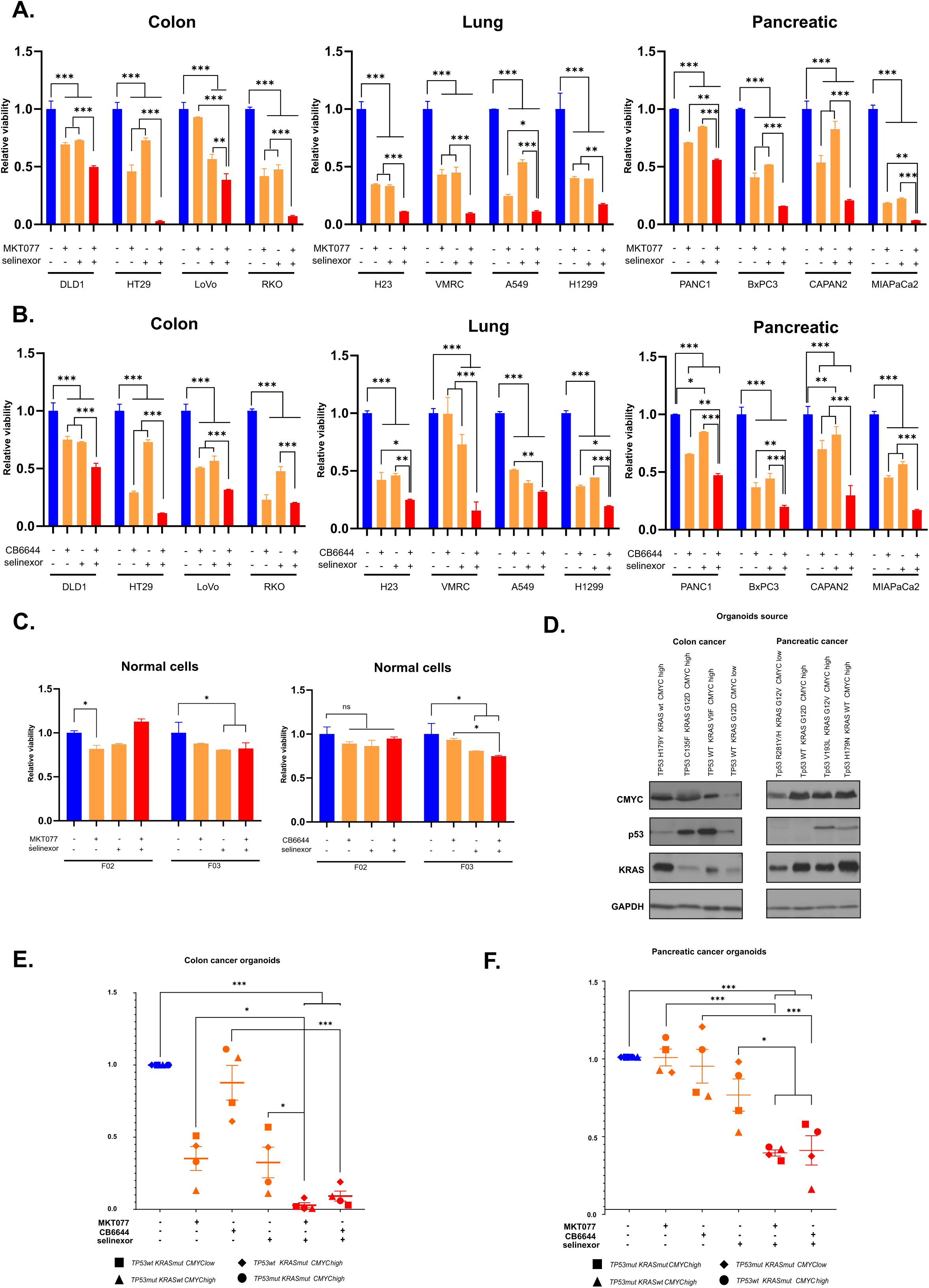
**A.** Impact of MKT077 (HSPA9 inhibitor) and selinexor (nuclear exportin 1 inhibitor), used as single agents and in combination, on viability of listed colon, lung and pancreatic cancer cell lines. **B.** Viability of listed colon, lung and pancreatic cancer cell lines treated with CB6644 (inhibitor of RUVBL1/2 ATPase activity), selinexor and combination of both inhibitors. **C.** Viability of normal fibroblasts upon treatment with indicated combinations of MKT077, selinexor and CB6644. Viability (A-C) was measured with ATPlite, 72 h post treatment with drug concentrations calculated individually for each cell line (based on IC_50_ values). Each bar represents mean of two replicates with SD. Data were analyzed with two-way ANOVA with Tukey’s correction, *p < 0,05, **p<0,01, ***p<0,001. **D.** Western blot indicating p53, KRAS and CMYC protein levels in the organoid cultures used in E-F, with the listed status of the three oncogenes in each culture. **E-F.** Viability of pancreatic and colon organoids, respectively, derived from tumor patient’s tissue, harboring listed mutations in *TP53*, *KRAS* and high/low CMYC level. 5 µM of MKT077, 2 µM of CB6644 and 2 µM of selinexor were used in combinations indicated in graph. Viability was measured 72 h after treatment using ATPlite assay. One-way ANOVA with Sidak’s correction was applied, *p < 0,05, **p<0,01, ***p<0,001.

To further confirm this observation we used *in vitro* model of heterogenous, patient-derived organoid cultures of pancreatic and colon cancers. The combination of selinexor with either MKT077 or CB6644 efficiently killed organoid cultures harboring mutations in *TP53*, and/or *KRAS* and/or high CMYC level (Fig. 3D-F). Significant viability decrease was accompanied by the loss of structure of treated organoids visible under a light microscope (Supplementary Fig. 3D). Obtained results altogether suggested that the combined blocking of exportin 1 and RUVBL1/2 complex, as well as inhibiting exportin 1 and HSPA9 may be tested in targeted therapeutic approach against cancers driven by mutant *TP53*, mutant *KRAS* and/or hyperactive *CMYC*.

### The druggable signature’s expression is associated with the presence of any of the activated trio of oncogenes in patient samples

To verify if the expression of the signature of the 3 genes *RUVBL1*, *HSPA9* and *XPO1* which encode proteins targeted in Fig. 3 is indeed associated with the presence of mutant p53, mutant KRAS or high level of CMYC, we interrogated cancer patient samples. We used locally collected samples from colon and pancreatic cancers patients’ tumors and TCGA datasets to validate the expression in multiple cancer types (Fig. 4). We tested the relative expression of the trio of individual genes in 24 samples of colon cancer (Fig. 4A) and 14 samples from pancreatic cancer (Fig. 4B). We stratified the samples using CMYC expression levels (“high” was above the mean of all samples in each cancer type) and presence of *TP53* or *KRAS* hotspot mutations in the samples (Supplementary Figure F, H). In case of each of the 3 tested genes the level of expression was on average significantly higher in samples containing any of the mutations and/or high CMYC overexpression (including samples with double conditions, indicated by horizontal lines linking the results in Fig. 4A and B), than the group of samples which contained none of the activated oncogenes (Fig. 4 and B). This indicated that high expression of *RUVBL1*, *HSPA9* and *XPO1* was significantly associated with the presence of any of the trio of activated oncogenes in the patient samples of colon and pancreatic cancers.

**Figure 4.**
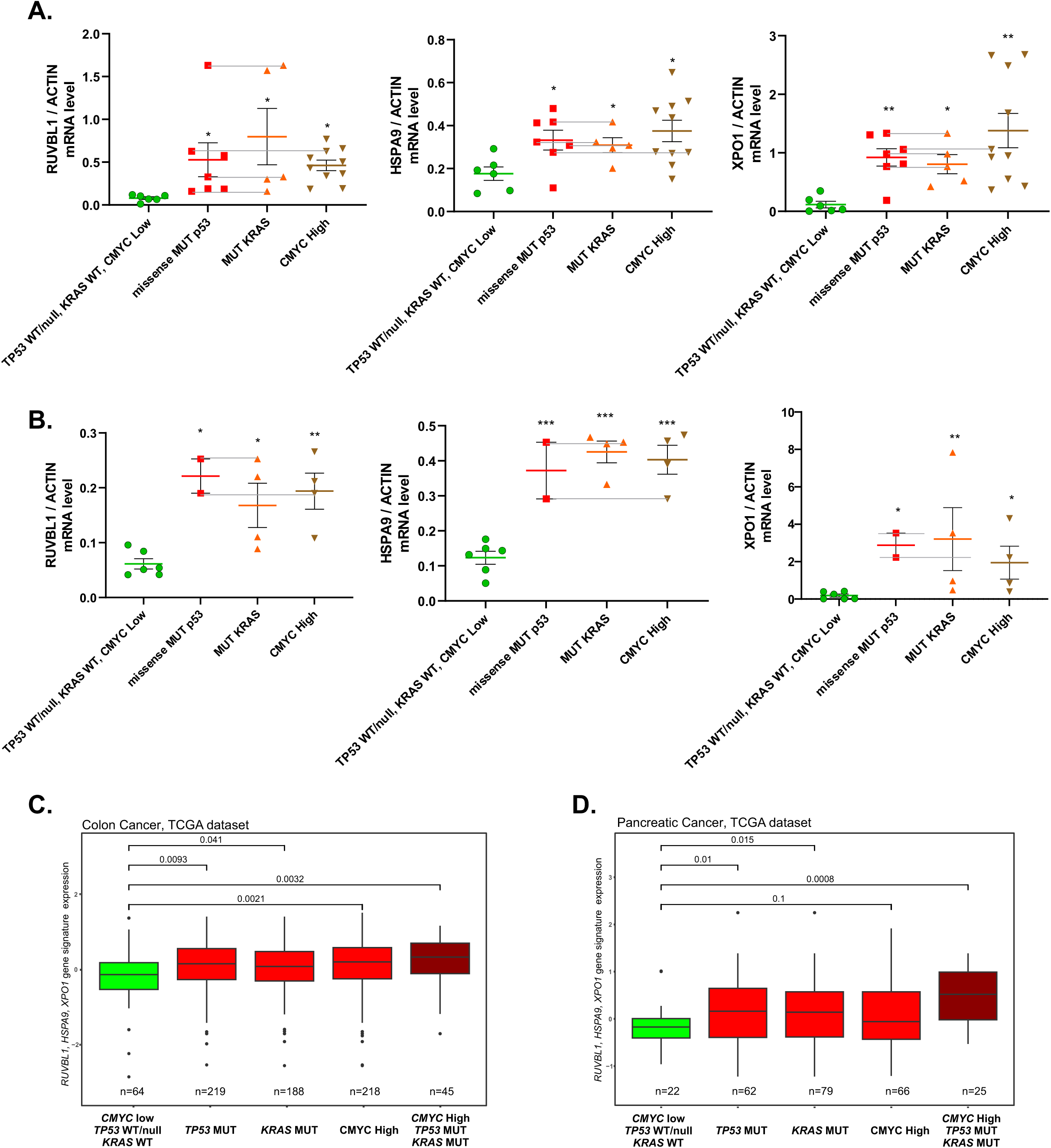
**A.** Expression of *RUVBL1, HSPA9* and *XPO1* genes was tested in 24 colon cancer samples stratified according to CMYC expression (samples above average of *CMYC* expression for all samples are considered “High”) and the presence of missense *TP53* and *KRAS* mutations. The samples with two coexisting oncogene activation conditions are linked with horizontal lines. **B.** The same expression analysis as in (A) for 14 pancreatic cancer samples. One-way ANOVA test with Durnett correction * p<0.05, ** p<0.01, *** p<0.001. **C.** Comparative expression analysis of a 3-gene signature consisting of *RUVBL1, HSPA9* and *XPO1* in TCGA-derived patient samples of colon cancer (mean value of three genes in each patient was used to calculate the sample distribution in the box plot), stratified according to the listed *TP53, KRAS* (point mutations only) and *CMYC* expression status. The sample was included in the “CMYC high” group if the *CMYC* gene expression was above the *CMYC* average expression level for all the patients in the graph. T-test was used to obtain the indicated p-vaules. **D.** As in (D) for TCGA-derived patient samples of pancreatic cancer.

To test if this association was present in larger populations of patients in publically available datasets we tested the signature consisting of *RUVBL1*, *HSPA9* and *XPO1* genes against those of TCGA-derived cancer patient datasets which contained sufficient *TP53* and *KRAS* mutations for statistically relevant analyses. The chi-square test result was instrumental in selecting the cancer types with the most significant prevalence of *TP53* and *KRAS* mutations (see Materials and Methods). The results for colon and pancreatic cancers are shown in Figure 4C and D respectively, while results for lung, uterine, bladder and stomach cancers are shown in Supplementary Figure 4 A-D. In all cases the presence of at least one activated oncogene of the studied *KRAS/TP53/CMYC* trio was sufficient to elevate the expression of the signature significantly (p<0.05) above the control group of patients (with *CMYC* expression below the average for all samples, and the WT *KRAS/TP53* status). In all cancer cases the co-presence of all three activated oncogenes was not resulting in the signature expression significantly higher than with a single active oncogene (Figure 4C and D; Supplementary Figure 4 A-D). This suggested that the oncogene activation is redundant rather than cooperative in increasing the signature expression.

We additionally tested if the 3-gene signature had a prognostic value in patient samples of the same cancer types as in the above-described expression analysis. In four out of six of the analyzed cancer types the signature’s high expression coincided with significantly worse patient survival, while in the cases of colon and stomach cancers this relation was reversed (Supplementary Fig. 4E). This indicated that while the signature represented usable treatment targets (Fig. 3) and its expression was consistently associated with the presence of the driver oncogenes (Fig. 4), its high level did not necessarily predict the worse survival of the patients in each studied cancer type.

### Expression of the targetable signature is redundantly and competitively controlled by the activated oncogenes

We then proceeded to asses if the activated oncogenes *CMYC*, mutant *KRAS* and *TP53* control the expression of *RUVBL1, HSPA9* and *XPO1* genes in a cooperative or in a redundant manner. We used uniform cellular background of K15 fibroblasts immortalized by introduction of hTERT ^17^ to overexpress via lentiviral transfection each of the oncogenes individually or in pairs (Supplementary Fig. 5A). We additionally silenced the introduced oncogenes to determine which of them remains in control of the target gene expression when the oncogenes are present in pairs. There were following observations in this experiment (Fig. 5 A-C for mRNA levels and Fig. 5D for protein levels): (i) each of the oncogenes - *CMYC*, mutant *KRAS* and mutant *TP53* – was individually able to activate each of the three target genes when compared to the expression level in the control K15 fibroblasts; (ii) there was no significant increase in expression of any of the targets when the oncogenes were co-overexpressed in pairs compared to overexpression of the individual oncogenes; (iii) the expression of the targets went down to the control levels upon silencing of the individually overexpressed oncogenes; (iv) when the oncogenes were overexpressed in pairs in most cases silencing of only one oncogene caused a significant decrease of the target gene expression. Interestingly, this last effect was did not include the same oncogenes for all the targets. While in the of co-presence of elevated CMYC and p53 R175H the expression *RUVBL1* and *XPO1* genes were dominantly under the CMYC control, the expression of *HSPA9* was dominantly controlled by the R175H p53 mutant (Fig. 5 A-C). All these results suggested that the expression of the analyzed oncogene targets are redundantly rather than cooperatively controlled by all the studied oncogenes, while the co-presence of the oncogenes usually lead to a dominance of one of them in inducing the target gene’s expression.

**Figure 5.**
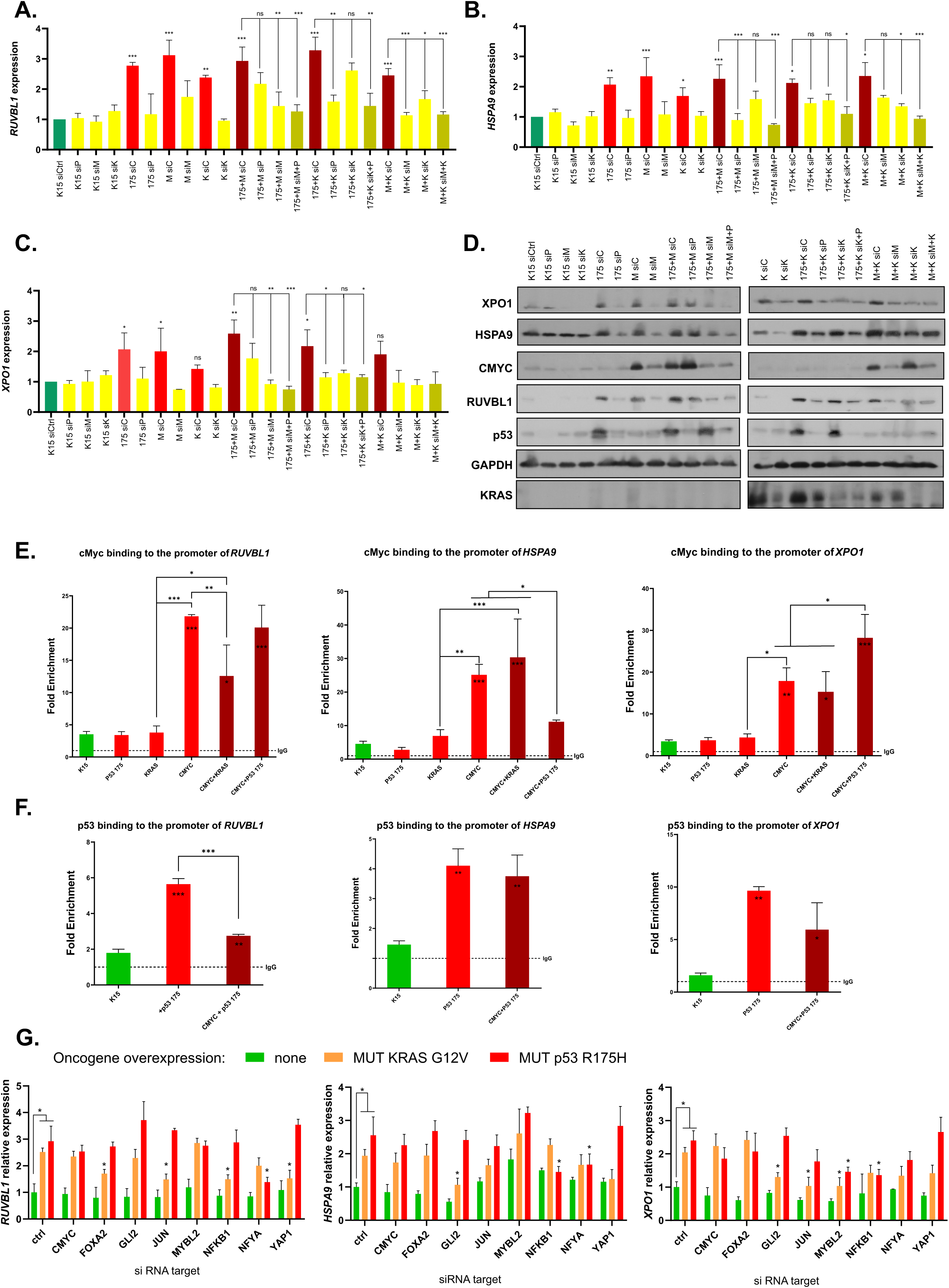
**A-C.** Relative mRNA expression levels of *RUVBL1, HSPA9* and *XPO1* genes (respectively) in K15 immortalized human fibroblasts upon indicated stable lentiviral vector-mediated overexpression and transient (48h) silencing of the oncogenes (overexpression: P – mutant p53 R175H, K – mutant KRAS, M – wt CMYC; “siX” - siRNA mediated silencing of the oncogene indicated by the letter). **D.** Western blot of the indicated proteins in K15 immortalized human fibroblasts upon the oncogene overexpression and silencing performed and described as in A-C. **F.** Chromatin immunoprecipitaion-derived qPCR result of the predicted CMYC-biding regions using anti-CMYC antibodies in the promoters of the indicated genes, performed in K15 immortalized human fibroblasts with listed stable oncogene overexpression or co-overexpression. The PCR results were normalized to the level of IgG unspecific antibody controls used for ChIP in parallel to specific antibodies in each oncogene overexpression setup. **G.** Chromatin immunoprecipitaion-derived qPCR result of the predicted CMYC-biding regions performed and normalized as in F. but using anti-p53 antibodies. **H.** Relative mRNA expression levels of *RUVBL1, HSPA9* and *XPO1* genes in K15 immortalized human fibroblasts upon indicated stable lentiviral vector-mediated overexpression of mutant *TP53* or *KRAS* and transient (48h) silencing of the listed candidate transcription co-factors. **A-H.** Means with SD are shown of 2-3 biological replicates (each a mean of 2 technical replicates), One-way ANOVA test with Durnett correction * p<0.05, ** p<0.01, *** p<0.001.

To investigate if a competition between the oncogenes for the dominance in controlling the target genes expression is reflected in binding of transcription factors to the target genes promoters we performed a chromatin immunoprecipitation (ChIP). CMYC and mutant p53 are known to bind to promoters (directly and indirectly, respectively) as well as to cooperate in this process ^12, 14^. We assessed their binding to promoter regions of *RUVBL1*, *HSPA9, XPO1* with the predicted CMYC biding sites in the K15 immortalized fibroblasts with introduced oncogenes (Materials and methods). We observed a significant increase of CMYC and R175H mutant p53 binding to all the promoters upon their overexpression in K15 fibroblasts, while the presence of mutant KRAS did not increase the presence of endogenous CMYC to the analyzed promoter regions. Notably, the co-expression of the oncogenes caused a significant increase of the CMYC binding only in the case of mutant p53 presence on the XPO1 promoter – suggesting cooperation of both oncoproteins. In all other cases there was either no effect of one oncogene on another, or we observed a significant decrease of the binding of one oncogene in presence of the other. This effect was e.g. visible in the case mutant p53 decreased binding to the *RUVBL1* and *XPO1* promoters upon overexpression of *CMYC*, while absent on the *HSPA9* promoter - where mutant p53 bound similarity with or without the high level of CMYC (Fig. 5F). This result was consistent with the earlier observation that when the oncogenes were co-present CMYC dominated the control of the *RUVBL1* and *XPO1* expression, while mutant p53 dominated over CMYC in the case of *HSPA9* expression (Fig. 5A-C). All these results suggested the existence of independence or competition between the oncoproteins, which eventually leads to redundant activation of the studied target genes.

The ChIP experiments (Fig. 5E-F) suggested that mutant KRAS and mutant p53 may regulate the analyzed target expression independently of CMYC. To find mediators of this activation we analyzed the promoters of *RUVBL1*, *HSPA9* and *XPO1* genes using several online-tools to find candidate transcription factors which may be induced by KRAS and mutant p53 (Materials and methods). Among the found factors we selected the top scoring common ones which were previously reported to be involved in KRAS or mutant p53 signaling (Supplementary table 6). Silencing of these selected transcription factors in the mutant *KRAS-* and mutant *TP53*-overexpressing K15 fibroblasts (Supplementary Fig. 5B) led to identification of significant dependencies, including GLI2 and CJUN which regulated expression of two genes each in the presence of overexpressed KRAS, while NFKB1 and NFYA regulated expression of two genes each in the presence of overexpressed mutant p53 (Fig. 5G). We additionally confirmed that indeed GLI2 and JUN bind the *RUVBL1*, *HSPA9* and *XPO1* promoters in the presence of the overexpressed KRAS mutant significantly stronger than in the control K15 fibroblasts (Supplementary Fig. 5C), while NFKB1 and NFYA are known to cooperate with mutant p53 ^2^.

These results indicate that each of the trio of the studied oncogenes has independent signaling routes that lead to the activation of the redundant targets, while in the presence of two activated oncogenes often one takes a dominant control of the target’s expression. As the example of *RUVBL1*, *HSPA9* and *XPO1* demonstrates, these redundantly controlled genes can be a source of novel druggable signatures of a broad specificity in various cancer and oncogene backgrounds.

### Redundancy between transcriptional programs of oncogenic CMYC, mutant KRAS and mutant p53 is a broad phenomenon in cancer cells

We sought to understand how broad is the phenomenon of redundancy in the transcriptional programs of oncogenic CMYC, mutant KRAS and mutant p53. First, we interrogated the transcriptional programs of the trio oncogenes determined earlier (Fig.1, Supplementary Table 2). For colon and lung cancer cell lines we overlapped mRNAs significantly changing levels (FDR<0.05) due to CRSPR-mediated downregulation oncoproteins as solo drivers, or in trios. In the case of each of the three oncogenes the overlaps allowed to determine control over which mRNAs remained specific to a given solo oncogene (non-redundant mRNAs), which was shared with other oncogenes (possibly redundant mRNAs, as the cooperative regulation is not excluded here), and which was taken over by other oncogenes when co-expressed (redundant mRNAs). In colon cancer cell lines shown in Fig. 6 A-C the transcriptional program controlled by CMYC was the most specific and the least redundant, as the majority of its target mRNAs were not co-controlled or taken over by mutant KRAS and p53 when co-expressed with CMYC (Fig. 6A). However, in the case of mutant p53 the majority of transcripts controlled by the solo-expressed oncogene were taken over by CMYC and/or mutant KRAS in the background of three oncogenes co-expression (Fig 6C). The redundancy of the mutant KRAS program was intermediate compared to CMYC and mutant p53 (Fig. 6B). In lung cancer cell lines CMYC retained even higher proportion of its solo-controlled mRNAs, while mutants of KRAS and p53 were strongly redundant to CMYC (Supplementary Fig. 6A-C). On average, in colon and lung cancer cell line setups tested, the transcriptional program of CMYC was the least redundant (6%), while the mutant p53 program was redundant in over 50% in favor of mutant KRAS and/or CMYC (Fig. 6D).

**Figure 6.**
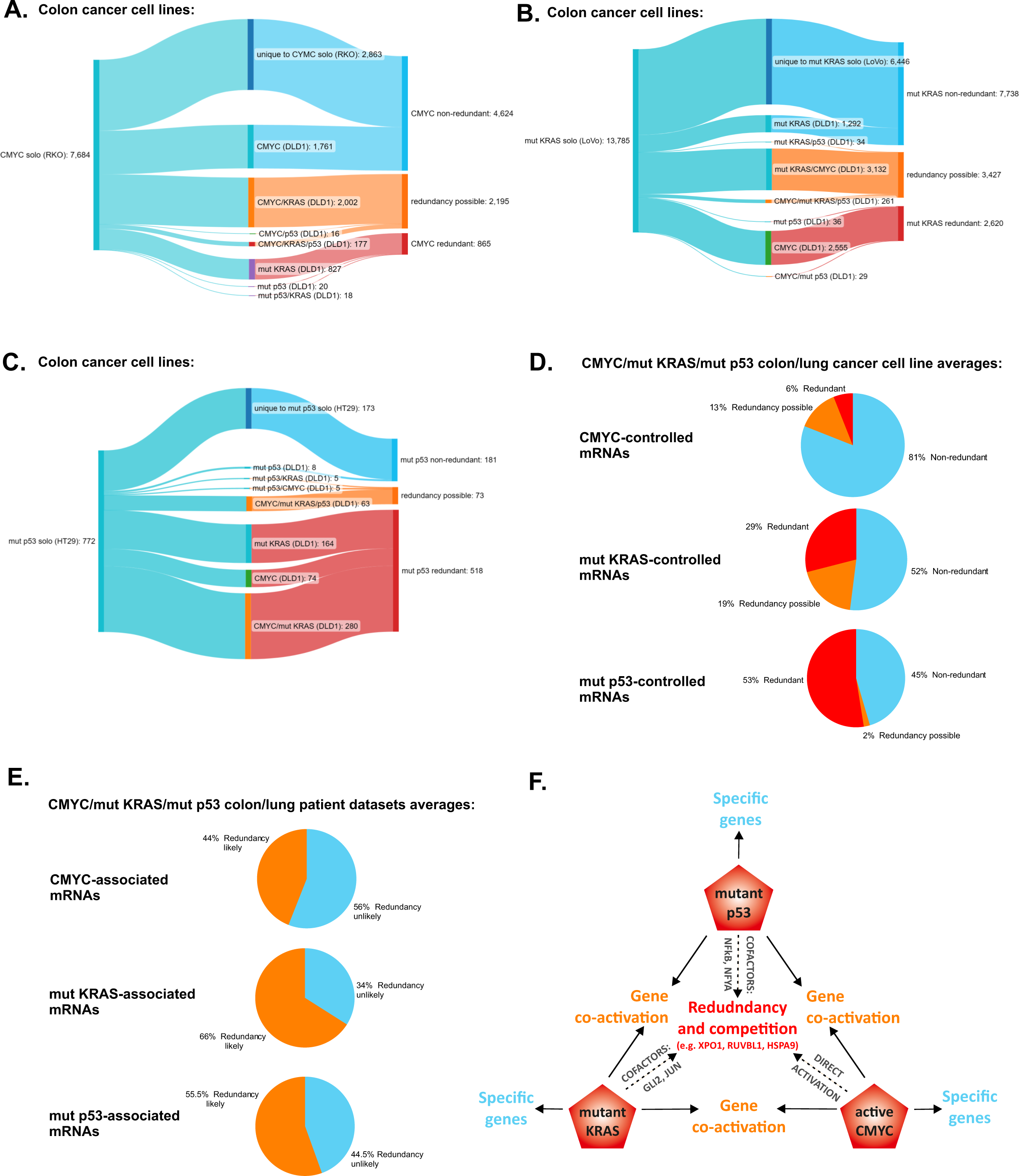
**A-C.** Ribbon charts showing gene pools dependent significantly (FDR<0.05; only coding mRNAs) on mutant *TP53*, mutant *KRAS* or hyperactive *CMYC* (respectively) in colon cancer cell lines with single activated oncogenes (left end) shared with cell line with three co-activated oncogenes (middle), resulting in: specificity to the single oncogene (non-redundant genes), sharing with the co-expressed oncogenes (redundancy possible genes), or take-over by the co-expressed oncogenes from the program of the single oncogene (redundant genes; right end). The data on differentially expressed genes is derived from the CRISPR-Cas9 experiment in Fig. 1A and Supplementary table 2. **D.** Percentages of non-redundant, possibly redundant and redundant genes for each indicated oncogene on average in colon (A-C) and lung (Supplementary Fig. 6 A-C) cancer cell lines. **E.** Percentages of genes with likely and unlikely redundancy associated with the listed oncogenes, on average in lung and colon cancer TCGA patient-derived expression datasets. The percentages were derived from gene pool flow analyses shown in Supplementary Fig. 6 D-E. **F.** Scheme depicting gene groups controlled by the studied oncogenes – genes specifically regulated by each oncoprotein, genes cooperatively controlled by the oncoproteins and redundant genes, which may also be a subject of the oncogene/oncoprotein competition – including the three gene signature described in the study.

We validated these results in a limited analysis against patient datasets from colon and lung cancers. This analysis was limited to association in contrast to earlier direct manipulation of the presence of the oncogenes, and thus we excluded combinations of tree oncogenes from the analysis, as it would be impossible to identify which genes remain controlled by the analyzed oncogene and which are taken over by the others. We overlapped mRNA programs significantly associated (p<0.05 for each gene) with the presence of the solo oncogenic CMYC, mutant KRAS, mutant p53 and the oncogenes other than the one under analysis to elucidate the scale of a possible redundancy of each oncogene (Supplementary Fig. 6 D, F for colon and lung cancers, respectively). Taking into account the average of results from colon and lung cancers the significantly regulated mRNAs associated with the presence of CMYC that were shared, and thus likely redundant, with other two oncogenes was 44%, while mutant p53 and mutant KRAS had 55.5% and 66% likely redundant genes, respectively (Fig. 6F). These results indicate that the oncogene redundancy range is dependent on the oncogene and it represents a phenomenon which may in selected cases involve the majority of the oncogene’s transcription driving potential.

## Discussion

The studies concerning omics of oncogene co-activation in neoplastic cells have been carried out in most cases using two experimental systems. One involves patient material-centered studies, which allow to associate activation of driver oncogenes with transcriptional/protein programs, phenotype features and help defining molecular subtypes of particular neoplasias ^18, 19, 20, 21^. Another experimental model involves introduction or activation of oncogenes in usually untransformed background *in vitro* or *in vivo* followed by functional phenotypic and omics analyses ^15, 22, 23^. Our study attempted to complement these methodologies with an approach involving CRSPR-mediated downregulation of the activated oncogenes in transformed cells *in vitro*. This allowed to directly compare the functional potential of each of the trio of major oncogenic drivers – mutant KRAS, mutant p53 and activated CMYC, solo and co-present with others – to control trascriptome and proteome in neoplastic cell addicted to the presence of the oncogenes. As expected from previous reports involving these oncogenes we found broad overlaps in the pathways driven by each of the oncogenes spanning across all three analyzed cancer types. Taking into account existing problems with direct targeting of each of the three studied oncogenes ^5, 8, 10^ and the universal character of the found overlaps we sought to find if the overlapping pathways contain usable targets for potential cancer treatment. This lead to finding of the three-protein signature of RUVBL1, HSPA9 and XPO1. Each of them individually was previously reported as a promising target in cancer ^24, 25, 26^, however synergistic pairwise drug combinations presented here exploit novel vulnerabilities against multiple cancer types, driven by the studied oncogenic trio.

From studies carried out so far we expected to find cooperative control of the target genes by mutant p53, mutant KRAS and CMYC. Multiple examples of how the oncogenes co-support expression of gene signatures were detailed in the Introduction and our earlier review ^2^. In contrast, results presented here, obtained using in-house patient-derived samples and public patient datasets, suggested that the control of *RUVBL1*, *HSPA9* and *XPO1* by the trio of the oncogenes was redundant, rather than cooperative. We examined the mechanism by introducing the oncogenes to immortalized, untransformed human fibroblasts and found that indeed each target gene is controlled by each oncogene in parallel, with cooperation found only in the case of mutant p53 and *CMYC* biding to the *XPO1* gene promoter, but not in the *XPO1* expression regulation. Moreover, we found that in most cases a specific oncogene dominates the control over a particular target, “switching off” the influence of the co-activated oncogene. In the case of the *RUVBL1* gene we observed a negative influence of mutant KRAS on biding of CMYC to the promoter, while CMYC negatively influenced mutant p53 binding to the *HSPA9* promoter. This “oncogene competition” on target promoters has been not been reported previously, while the existence of this mechanism is likely, taking to account studies on other competing oncogene functions ^27^ or competition of transcription factors out of the oncogenic context ^28, 29, 30^.

In attempt to assess the prevalence of the oncogene redundancy we found that the proportion of genes whose control is carried over from one oncogene to two others depends on the oncogene – CMYC being the most redundancy-resistant and mutant p53 being redundant on average in over 50% of its transcriptional program in lung and colon cell lines. This result was reflected by the oncogene-specific programs in lung and colon cancer patient datasets, although, unlike the direct functional experiment, this analysis was limited to association.

The concept of more and less “powerful” oncogenes, with mutant p53 largely dominated by the presence MYC/KRAS, could be explained by larger gene numbers and stronger expression changes induced by the latter (Fig. 1), leading to stochastic gene co-regulation. However, the case of mutant KRAS significantly affecting expression of a larger number of mRNAs in colon cancer cells and still being more redundant to CMYC than the other way around (Fig. 6A, B), suggests that the mechanism is more complex than just the domination of a broader expression regulator. The scale of the redundancy, reaching over 50% of mutant p53-driven transcripts, is potentially critical for the oncogene function and targeting by drugs. It is also likely reflected in a controversy around p53 mutants as driver oncoproteins. In the majority of studies mutant p53 oncogenic gain-of-function has been found and explained ^2, 31^. However, there are cancer models where mutant p53 does not show a clear gain-of–function, exerting only dominant-negative effect and loss of oncosuppressive function ^32, 33, 34^. Such cases have been hypothesized to be dependent on a tissue/molecular context ^31^. Based on results shown here we propose that other activated oncogenes are one important part of the background context, defining the range of influence of mutant p53 on cancer cells.

It remains to be evaluated to what extent the “race for power” between the oncogenes involves the oncogene competition and includes other drivers than the trio described here. Nevertheless, the extensive oncogene redundancy comprises a warning of possible resistances arising from direct targeting of the oncogenic drivers - as already reported in FGFR inhibitor clinical trials ^35^. This mandates a search for targetable signatures in cancer, such as RUVBL1, HSPA9 and XPO1 shown here, whose redundancy is limited by directly non-overlapping roles in cell metabolism.

## Funding

The research was financed by National Science Center, Poland, Sonata Bis grant no. 2017/26/E/NZ5/00663 and PTO/Servier Oncology Grant (1^st^ edition) to D.W., and MMRC PAS internal grant FBW-03/2021 to M.G.

## Acknowledgements

We thank prof. Harm Kamipinga (University of Groningen, The Netherlands) for providing K15 immortalized fibroblasts and dr Damian Graczyk (Institute of Biochemistry and Biophysics PAS, Warsaw, Poland) for assistance with colon cancer cell lines. We thank all cancer patients from hospitals in Warsaw, Poland, who provided consent to use their tissue samples in the manuscript as a part of the “Multi-onko-mapa” study.

## Author contributions

Conceptualization, D.W. and M.Grz.; Methodology, D.W. M.Grz., M.Gro., A.J., G.C., S.P., and J.R.W,; Investigation, D.W. M.Grz., M.Gro., A.J., J.R.W; Data curation, D.W., J.R.W. and A.J., Writing – Original Draft, D.W and M.Grz.; Writing – Review & Editing, D.W. and M.Grz.; Funding Acquisition, D.W. and M.Grz.; Resources, M.N-N., M.K., W.K., T.O. and M.L.; Supervision, D.W.

## Materials and methods

### Cell lines

RKO (RRID:CVCL_0504), HT29 (RRID:CVCL_A8EZ), PANC-1 (RRID:CVCL_048), MIA PaCa-2 (RRID:CVCL_0428), NCI-H23 (RRID:CVCL_1547), NCI-H1299 (RRID:CVCL_0060), A-549 (RRID:CVCL_0023) DLD-1 (RRID:CVCL_0248), Capan-2 (RRID:CVCL_0026) and BxPC-3 (RRID:CVCL_0186) cell lines were acquired from the American Type Culture Collection (ATCC). VMRC-LCD (RRID:CVCL_1787) cell line was purchased from Japanese Collection of Research Bioresources Cell Bank (JCRB). LoVo (RRID:CVCL_0399) cell line was acquired from the European Collection of Authenticated Cell Cultures (ECACC) repository. For all the experiments only low passage numbers (below 15 post acquisition) of cell lines were used. The cells had routinely excluded presence of *Mycoplasma* sp. by qPCR.

A-549, VMRC-LCD, RKO, LoVo, HT29, PANC-1 and MIA PaCa-2 cell lines were cultured in DMEM medium (Gibco) supplemented with 10% FBS (Gibco) and 1% Pen Strep Antibiotics (Gibco). NCI-H23, NCI-H1299, DLD-1, Capan-2 and BxPC-3 cell lines were cultured in RPMI medium (Gibco) supplemented with 10% FBS (Gibco) and 1% Pen Strep Antibiotics (Gibco). Human primary fibroblasts (F02 and F03) were obtained from skin biopsies of healthy subjects based on bioethics committee approval (No 108/2017 and 203/2020) of the Central Clinical Hospital of Ministry of Interior and Administration in Warsaw ^36^. Human h-TERT immortalized K15 fibroblast were a kind gift from Prof. Harm Kampinga (Univ. of Groningen) ^17^. All fibroblasts were cultured in DMEM medium (Gibco) supplemented with 10% FBS (Gibco) and 1% Pen Strep Antibiotics (Gibco).

### Lentivirus production and transfection

Lentiviruses for protein overexpression were produced by transient plasmid transfection of HEK293T cells (ATCC). Briefly, HEK293T cells were seeded in 10-cm dishes and the following day were transfected with two second-generation packaging plasmids - 5 µg pMD2.G and 10 µg psPAX2 (gift from Prof. Giannino Del Sal, ICGEB Trieste, Italy) and 15 µg of target lentiviral vectors carrying: Cas9 cds (a gift from Dr Magdalena Winiarska, Warsaw Medical University/MMRC PAS), CMYC cds (Addgene #46970), KRAS4B G12V cds (Addgene #35633), p53 R175H or R273H cds (a gift from Dr Maciej Olszewski, IIMCB, Warsaw). 90 µg of PEI (Sigma Aldrich) was used for transfection. After 48 hours the medium was collected, filtered through 0.45µm PVDF filters (Millipore), enriched with 10% of FBS and 1% Polybrene (Sigma Aldrich) and used for target cells infection. After 48-72h the medium containing viruses was replaced with fresh media with 1 μg/mL Puromycin or 1 μg/mL Hygromycin B Gold (InvivoGen), used for selection.

### CRISPR/Cas9-mediated oncogene downregulation

The gRNAs, used for CRISP/Cas9-mediated *KRAS*, *CMYC* and mutant *TP53* silencing, was annealed from two components: trans-activating crRNA (A35507, Invitrogen) and crRNA targeting specific gene (A35509, Invitrogen) or negative control (A35519, Invitrogen), according to manufacturer’s protocol. For *KRAS*, *CMYC* and *TP53* targeting we used a mix of two crRNA in 1:1 proportion (CRISPR577256_CR and CRISPR721685_CR for KRAS, CRISPR634316_CR and CRISPR634324_CR for CMYC, CRISPR718498_CR and CRISPR718512_CR for *TP53*), in order to improve the downregulation efficiency. Cancer cell lines with z stable Cas9 overexpression introduced by lentiviral infection were seeded in 6-cm dishes and transfected with gRNA mixes using Lipofectamine MessengerMax (Invitrogen), following manufacturer’s instructions. After 48 hours cells were harvested for oncoprotein level validation, RNA-sequencing and proteomics analysis.

### Western blot and antibodies

The effectiveness of CRISPR/Cas9 treatment or oncogene overexpression/silencing was assessed by western blot. Cells were collected and lysed in NP40 buffer (150 mM NaCl, 1% NP-40, 50 mM Tris-HCl, pH 8.0) with HALT protease inhibitor (Thermo Fisher Scientific). After 10 min incubation in Laemmli Sample Buffer in 95°C, samples were run using 10% SDS-PAGE gel and transferred to nitrocellulose membrane (Merck). Membranes were blocked with 5% fat-free milk in TBS-Tween20 0.1% and incubated overnight with primary antibodies listed in the Supplementary Table 7.

### Proteomics analysis

Cells after CRISPR/Cas9 treatment (in 3 biological replicates of control and each oncogene downregulation) were lysed in buffer (50 mM Tris-HCl, pH 7.8) containing 1% SDS and 0.1 M dithiothreitol and sonicated (Diagenode Bioruptor Plus). Number of sonication cycles was established based on a visual sample clarity assessment. WF-assay was applied in order to calculate protein and peptide concentration ^37^. Multi-Enzyme Digestion Filter Aided Sample Preparation (MED FASP) protocol ^38^ with minor modifications ^39^ was applied for lysate processing. Briefly, endoproteinase LysC was used for overnight protein digestion, followed by 3 h incubation with trypsin. 0.5 mg of peptides were separated on a reverse phase C18 column and analyzed on QExactive HF Mass Spectrometer (Thermo-Fisher Scientific) as described previously ^40^. MaxQuanf software (https://maxquant.net/maxquant/) was used for spectra search and total protein approach using the raw protein intensities was applied for protein concentration calculation. Perseus software (https://maxquant.net/perseus/) was used to perform differential analysis, t-tests and assess p-value support of differences between protein concentrations in distinct experimental conditions as well as perform hierarchical clustering.

### RNA-sequencing

Total RNA was extracted from cell lines in 3 biological replicates after each CRISPR/Cas9 oncogene downregulation using QIAzol (Qiagen) following the manufacturer’s instructions. Quality and quantity of obtained RNA was analyzed using NanoDrop (Thermo Fisher Scientific) and Experion RNA analyzer (Biorad). After additional quality control (Qubit RNA, Thermo Fisher) and Bioanalyzer RNA (AGgilent) and libraries preparation (KAPA RNA HyperPrep with RiboErase, HMR), samples sequencing (NovaSeq6000; 100M reads, 2x100bp) and preliminary data quality check was performed by CeGat GmbH (Germany).

Quality assessment of the raw data including filtering and trimming was carried out using FastQC_v.0.11.9 (http://www.bioinformatics.babraham.ac.uk/projects/fastqc/) and the data was further processed using tool Trimmomatic_v.0.36 (http://www.usadellab.org/cms/?page=trimmomatic) for removal of low quality reads (phred-score ≤ 20 and length 30 bp, singletons discarded, reads with ambiguous bases ‘N’), trimming of bases from 5’/3’ end and adaptor sequences. Refined and filtered reads were further mapped to the reference human genome by Bowtie2 – used for indexing (http://bowtie-bio.sourceforge.net/bowtie2/), and HISAT2 – used for mapping (http://daehwankimlab.github.io/hisat2/). The successfully mapped reads were further processed to FeaturesCount R package to calculate the abundance of each transcript and extract quality score of mapped reads. Good quality scores were saved in a matrix form and were used as an input for determination of differentially expressed genes between control and oncogene dowregulation conditions by using DESeq2. The Relative Log Expression (RLE) method was used in DESeq2 to calculate normalization factors. Benjamini-Hochberg false discovery rate, FDR<0.05 was considered as statistically significant parameter of significantly differentially expressed genes. The count matrix genes were annotated with Ensembl BioMart to extract protein coding genes, which were used to generate principal component analysis (PCA) plot. Perseus software (https://maxquant.net/perseus/) was used to perform hierarchical clustering of the DEA results.

### Pathway analysis and target gene extraction

Proteins and mRNAs significantly changing levels (p<0.05 and FDR<0.05, respectively) in the differential analysis of proteomes and transcriptomics for each oncogene were fused in to signatures and filtered for duplicates. Such signatures were used in ClueGO ver. 2.5.8 (42) plug-in in Cytoscape ver. 3.8.2 (www.cytoscape.org) to associate proteins with molecular pathways. ClueGO settings were: All_Experimental evidence, GO Molecular Pathways/KEGG/WikiPathways ontologies, network specificity slider half way between Medium and Detailed settings, show only pathways with p<0.05. Analyses performed for the separate oncogenes - were exported into tables and overlapped (Supplementary Table 3) to determine pathways specific and common to the signatures. Presence of each gene associated with the pathways common to all three oncogenes was then validated in each of the proteomics and teranscriptomics significant DGE results from each cell line, and the genes with highest counts were selected to represent the pathways in the heat map in Fig. 1F.

### Gene expression and survival analysis in patient datasets

The analysis was conducted using the patient gene expression data from TCGA datasets for cancer types indicated in the figures, such as lung, colon, pancreatic, stomach, uterine and bladder urothelial cancer. Patients were stratified based on CMYC expression levels “high” and “low” as below or above mean level in each dataset, and the presence of *TP53* or *KRAS* point mutations. For Figure 4 a comparative expression analysis of the 3-gene signature (*RUVBL1, HSPA9, XPO1*) was conducted, categorizing patients into specific groups such as *CMYC* high only, *KRAS* mutation only, *TP53* missense mutation, then *CMYC* high + *TP53* mut + *KRAS* mutation group *CMYC* low + *TP53* WT/null + *KRAS* WT as a control group. The T-test was employed to calculate significance for each cancer type separately and box plots were generated using ggplot R packages for each cancer type. For analysis in Figure 6 similarity stratified patient datasets were used to compare expression levels of all mRNA available in each dataset for a given expression/mutation set with the rest of the set. T-test with FDR Benjamini-Hohberg correction result cut-off of <0.001 was used to determine mRNAs whose significant change is associated with the presence of particular expression/mutation set. Such mRNA lists were used for overlaps shown in ribbon plots in Figure 6.

Survival analysis in Supplementary Figure 4 was done using Kaplan-Meier plots with log-rank tests, categorizing patients into low and high expression groups for the 3-gene signature (*RUVBL1, HSPA9, XPO1*). The log-rank test assessed significant differences in survival rates, while hazard ratios estimated relative risks. The findings aimed to elucidate the relationship between the expression levels of signature genes and patients’ survival across diverse cancer types. This analysis was conducted using the Surv function in R, and plots were constructed using the ggplot packages.

### Promoter binding candidates selection

Three genes form the studied signature were each separately input to four online tools assisting determination of transcription factors binding gene promoters: <u>ChEA3 (maayanlab.cloud)</u>, <u>EPD The</u> <u>Eukaryotic Promoter Database</u> (expasy.org), <u>** CiiiDER **</u> and <u>ChIP-Atlas</u>. Top hits were overlapped between the genes and the transcription factors suggested by each tool for at least two genes were further considered. The final choice (Fig. 5, Supplementary table 6) was based on cross-checking the top hits with the literature data confirming functional link with KRAS and/or mutant p53.

### siRNA sinencing and mini-screen

For the mini-screen the cells were plated in 96-well plates and transfected with 20 nM of pre-designed siRNAs (purchased from Horizon Discovery – Dharmacon, listed in Supplementary Table 4) with the use of Lipofectamine RNAiMAX (Invitrogen), according to manufacturer’s instructions. 48 hours after transfection 1% resazurin (Sigma) was added to each well and 2 h later viability was measured.

For experiments with the use of siRNAs in K15 fibrobalsts, cells were seeded in 6-12 well plates, and transfected with indicated siRNAs (listed in Supplementary table 7) with the use of Lipofectamine RNAiMAX (Invitrogen), according to manufacturer’s instructions using Lipofectamine RNAiMAX (Invitrogen). 48h hours after transfection cells were harvested and further processed.

### Drug tests

CB-6644, MKT-077 (both purchased from MedChemExpress) and selinexor (KPT-330, Selleck Chemicals) were dissolved in DMSO and used in concentrations calculated individually for each cell line, as described in Results section. Cancer cell lines as well as normal fibroblasts were seeded in 96-well plates with clear bottom. After 24h medium was replaced for fresh one with single drug or drugs combination. 72h hours later viability of cells was measured using ATPlite One Step Reagent (Perkin Elmer).

### Total RNA extraction from cell lines and patient’s samples

Total RNA from cell lines was extracted using RNA Extracol (EURx, Poland), following standard phenol-chloroform RNA isolation protocol. Patient’s frozen samples were homogenized, incubated in RNA Extracol with RNA extraction beads (Diagenode), sonicated using Bioruptor Plus (Diagenode), and further processed according to standard phenol-chloroform RNA isolation protocol. RNA quantity and quality was assessed utilizing NanoDrop spectrophotometer (Thermo Fisher Scientific).

### RT-qPCR

500 ng of total RNA was reverse-transcribed using NG dART RT kit (EURx, Poland), according to manufacturer’s protocol. Sensitive RT HS-PCR Mix SYBR (A&A Biotechnology, Poland) reagents were used for qPCR on One Step Plus Real-Time PCR System (Applied Biosystems) and CFX Maestro (Biorad). Primers used for qPCR are listed in Supplementary Table 7.

### Human frozen tissue samples

Sample collection and further laboratory experimental procedures were performed based on ethical committee approvals: No 109/2016 (with updates) of National Medical Institute of Ministry of Interior and Administration in Warsaw and No 55/2023 of National Insitutte of Oncology in Warsaw. Written consents for research use of collected tissues were obtained from all patients. Altogether 28 samples of colon cancer and 18 samples of pancreatic cancer were collected from patients undergoing surgical treatments (Figures 4S and 5S). Samples, after surgical resection, were subjected to preliminary histopathological assessment and further storage in culturing medium at 4°C for organoid culture establishing or in liquid nitrogen for further DNA, RNA and protein extraction. For tumor type and grade classification according to WHO guidelines, tissues were fixed in 10% buffered formalin, embedded in paraffin, microtome cut into 4 mm thick sections and HE (haemotoxylin/eosin) stained.

### Human colon cancer organoid cultures

Colon cancer and normal colon tissues were transported at 4°C in culturing medium w/o growth factors and processed within 18 h from resections. Protocol from ^41^ with small modifications was used to generate colon organoids. First, tissues were washed 10 times with ice-cold PBS, then minced using surgical scalpel. Minced tissues were incubated in digestion medium (Collagenase type II 5 mg/mL [GIBCO], Dispase 5 mg/mL [GIBCO], Y-27632 10.5 μM [Sigma], DNase I 10 μg/mL [Sigma], Advanced DMEM/F12 [GIBCO], HEPES 10 mM pH 7.5 [Invitrogen], GlutaMAX Supplement 1× [Invitrogen], Primocin 100 ug/mL [InvivoGen], Bovine Serum Albumin 0.1% [Sigma]) for 40 min at 37°C with rotation. Remaining undigested tissue fragments were allowed to settle to the bottom of the tube for 1 min, then the supernatant was collected in a 15 mL Falcon tube, centrifuged at 300 RCF for 5 min at 4°C. The cell pellet was embedded in a growth factor reduced Matrigel (Corning) or Cultrex Basement Membrane Extract type 2 (R&Dsystems). After 20 min incubation at 37°C, matrigel domes were covered with the culturing medium (Advanced DMEM/F12 [GIBCO], HEPES 10 mM pH 7.5 [Invitrogen], GlutaMAX Supplement 1× [Invitrogen], 10% R-spondin-1 conditioned medium (Ootani et al., 2009), 50% Wnt-3A conditioned medium (Sato et al., 2011), N-acetylcysteine 1.25 mM [Sigma], Nicotinamide 10 mM [Sigma], B27 supplement 1× [GIBCO], Primocine 100 ug/mL [InvivoGen], murine Noggin 100 ng/mL [Peprotech/ThermoFisher], human EGF 50 ng/mL and human FGF-10 100 ng/mL [Peprotech/ThermoFisher], A83-01 500 nM [Tocris], Y-27632 10.5 uM [Sigma], SB202190 3 uM [Sigma]).

### Human pancreatic cancer organoid cultures

Pancreatic organoids were generated in a similar way to colon organoids based on ^42^, with addition of human Gastrin I 10 nM [Tocris] and no SB202190 in the culturing medium. Prostaglandin E2 10 nM [Tocris] was added only to medium for normal tissue organoids.

### Drug sensitivity assays in organoids

For organoids 96-well plates with clear bottom and white walls were used for testing. 33 μL of Matrigel or Basement Membrane Extract type 2 were added to each well. Plates were then centrifuged at 1000 RCF for 1 min and placed in a 37°C, 5% CO2 incubator for 30 min. To each well a 100ul suspension of approximately 500 organoids in the culturing medium were added. Organoids were left for 24 h in a 37°C, 5% CO2 incubator. Afterwards the drugs were added in the culturing medium at concentrations indicated in the figures. The organoids were incubated with drugs for 72 h, then cell viability was measured using ATPlite Luminescence Assay (PerkinElmer).

### Chromatin immunoprecipitation

Chromatin of live cells was cross-linked with 1% formaldehyde for 20min at room temperature. Cross-linking was stopped by incubating cells with 125mM Glycine/PBS for 5min. Cells were scraped in PBS, centrifuged and then lysed with Lysis Buffer (50mM HEPES pH 8, 140mM NaCl, 1mM EDTA, 10% glycerol, 0.5% NP-40 and 0.25% Triton X-100). Cell pellet was washed with Wash Buffer (10mM Tris-HCl pH 7.5, 200mM NaCl, 1mM EDTA pH 8). Chromatin was sonicated in Shearing Buffer (0.1% SDS, 1mM EDTA and 10mM Tris pH 7.5) using Bioruptor sonicator (Diagenode; high power setting) to obtain fragments of about 500-800 bp. The buffer was then supplemented to obtain RIPA-100 Buffer (20mM Tris-HCl ph 7.5, 100mM NaCl, 1mM EDTA, 0.5% NP-40, 0.5% Na-Deoxycholate, 0.1% SDS). Samples were precleared for 1h at 4°C with the addition of Protein A/G PLUS-Agarose (Santa Cruz Biotechnologies). Chromatin was immunoprecipitated overnight at 4°C using antibodies: p53 DO-1 (Santa Cruz sc-126); cMyc/NMyc (D3N8F Cell Signaling); Gli-2 (Santa Cruz sc-271786); c-Jun (Santa Cruz sc-74543); NF-YA (Santa Cruz sc-17753); NFKB p50 (Santa Cruz sc-8414). Rabbit serum IgGs (Novus) were used as a negative control. Protein A/G PLUS-Agarose was added to chromatin immunoprecipitates to recover DNA-protein complexes followed by washing the complexes with RIPA-100 Buffer, then RIPA-250 Buffer (10ml of RIPA-100 supplemented with 300μl NaCl 5M), LiCl Buffer (10mM Tris-HCl pH 7.5, 1mM EDTA, 250mM LiCl, 0.5% Na-Deoxycholate, 0.5% NP-40) and finally with TE Buffer (1mM EDTA pH 8, 10mM Tris-HCl pH 8). DNA-protein complexes were then treated with RNase A for 30min at 37°C in TE Buffer. Next, Proteinase K Solution was added (1% SDS, 200mM NaCl, 300μg/ml Proteinase K) and samples were incubated at 68°C overnight. All the buffers were supplemented with protease inhibitors. Coimmunoprecipitated DNA was extracted using. Coimmunoprecipitated DNA was analyzed using real-time PCR (Biorad CFX Opus 96 Real-Time PCR System) and SYBR Green Sensitive RT HS-PCR Mix SYBR (A&A Biotechnology). Promoter occupancy was calculated using Fold Enrichment Method (2−ΔΔCt method). PCR primers for expected CMYC biding sites were designed using ChIP-Atlas (https://chip-atlas.org/) and are listed in the Supplementary Table 7.

### Statistical analysis

GraphPad Prism 8.0.2 was used for statistical analysis and data visualization. Data are presented as mean ± standard deviation (SD) or standard error of the mean (SEM). Number of replicates as well as applied statistical test are described in each figure legend.

## Notes

### Competing Interest Statement

The authors have declared no competing interest.

