## Supplementary Figures and Legends for "A common druggable signature of oncogenic CMYC, mutant KRAS and mutant p53 reveals functional redundancy and competition of the oncogenes in cancer"

### Supplementary Figure 1

## A.

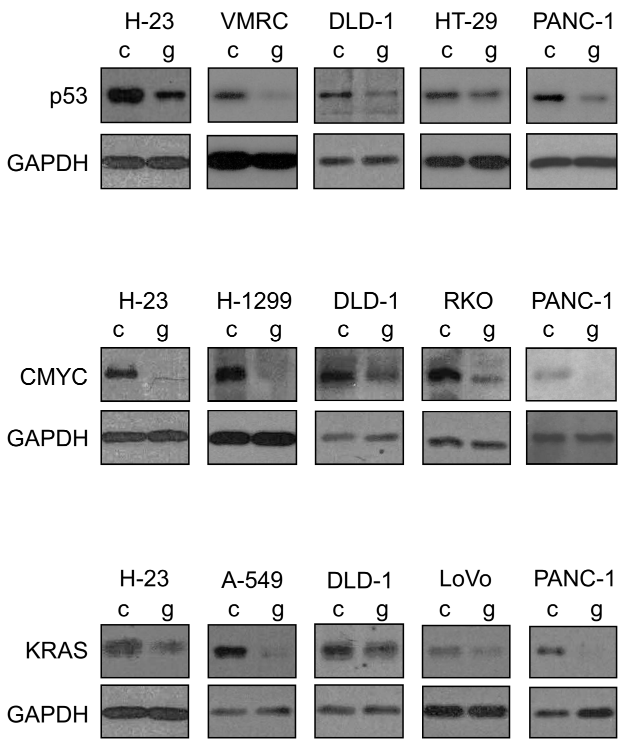


## B.

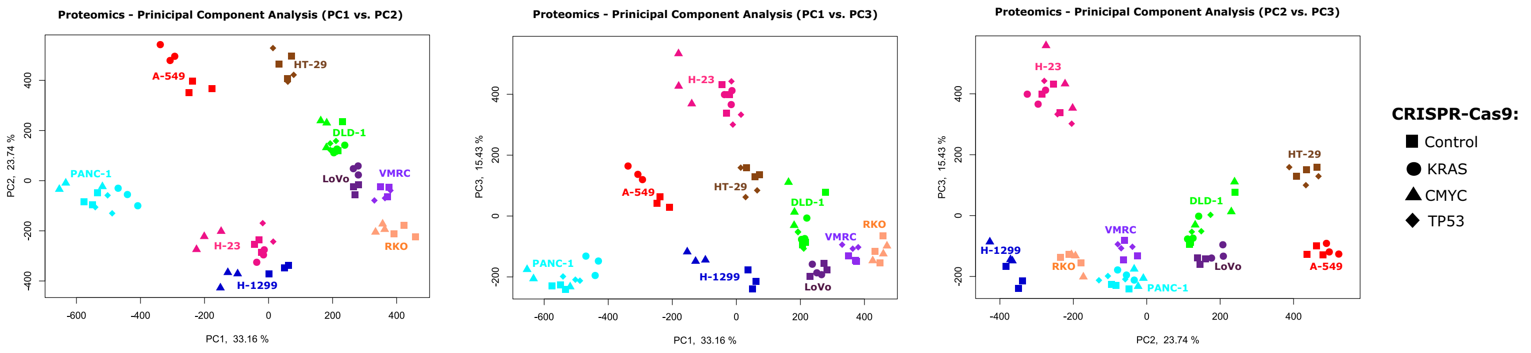


## C.

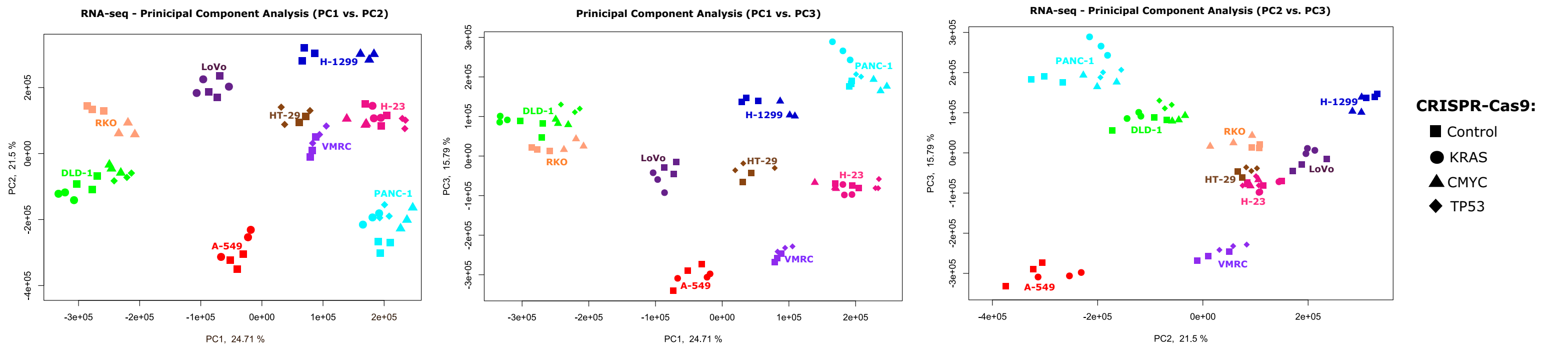

**Supplementary Figure 1** **A.** Western blots showing representative samples of control and oncogene-targeting gRNA in CRISPR-Cas9-mediated downregulation of indicated oncoproteins, whose large-scale analysis is shown in Figure 1. Analyzed cell line is indicated above each WB. **B.** Principal component analysis (PCA) of proteomics samples described in Figure 1. Cell lines are marked by colors and names, while gRNAs used in CRISPR-Cas9 experiments are indicated by symbol shapes. **C.** PCA for mRNAs resulting from RNA-sequencing described in Figure 1 done and shown as in B.

### Supplementary Figure 2

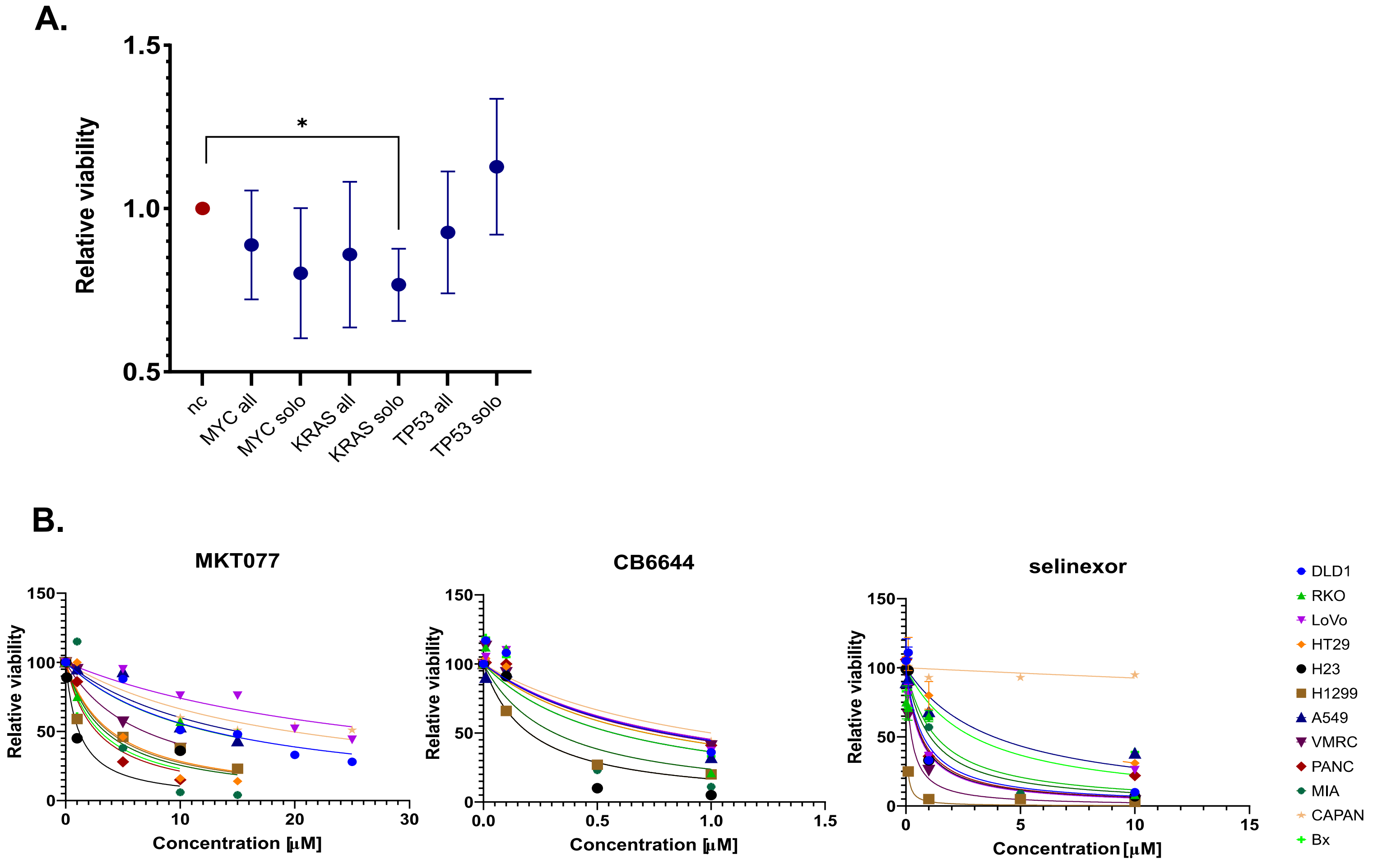

**Supplementary Figure 2** **A.** Average viability of cell lines transfected with siRNAs targeting indicated oncogenes, carrying either single oncogene or all three. Resazurin assay was used for viability measurement 48h post siRNA transfection. Shown are means of 3 cell lines' results for each oncogene, normalized to control siRNA (nc) viability level, and analyzed with one-way ANOVA (uncorrected Fisher's LSD) versus the siRNA control result (* $p<0.05$). **B.** Titration of indicated inhibitor concentrations in listed cancer cell lines shown as function of their viability measured 72 h post treatment with the use of ATPlite. The fitted inhibition curves were used to calculate IC50 values listed in Supplementary Table 5.

#### Supplementary Figure 3

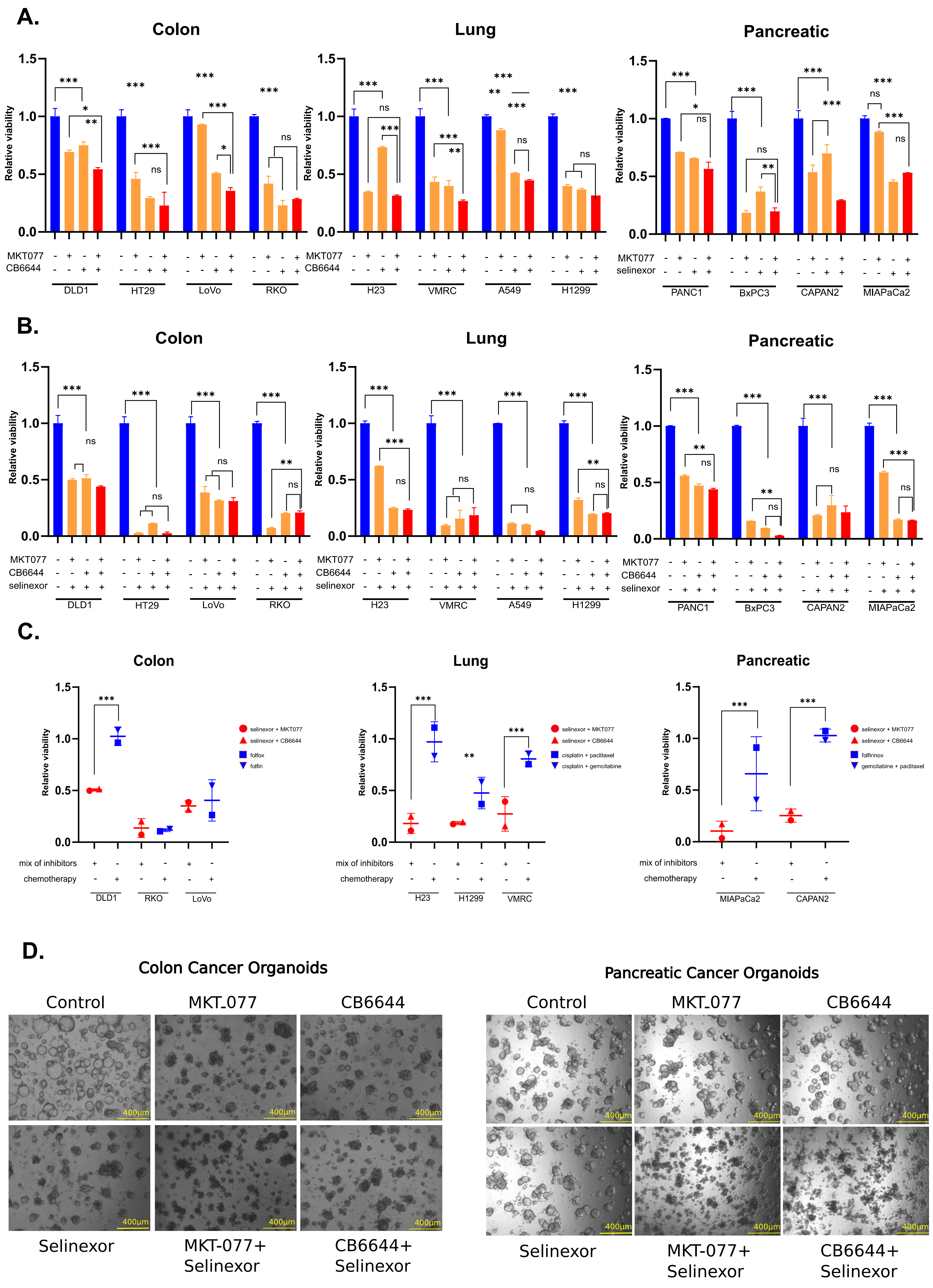

**Supplementary Figure 3** **A.** Viability of colon, lung and pancreatic cancer cell lines treated with CB6644, MKT077 and combination of both inhibitors. **B.** Impact of MKT077, CB6644 and selinexor mixture on viability of colon, lung and pancreatic cancer cell lines. Viability in (A-B) was measured with ATPlite reagent, 72 h post treatment with drug concentrations calculated individually for each cell line (based on IC50 values in Supplementary Table 5). Each bar represents mean of two replicates with SD. Data were analyzed with two-way ANOVA with Tukey's correction, *p < 0,05, **p<0,01, ***p<0,001. **C.** Comparison of viability of indicated cell lines with most effective inhibitor combinations introduced in this study (selinexor+MKT077 and selinexor+CB6644 – mean with SEM is shown) with two listed standard chemotherpautic protocols for each cancer type (mean of the viability results with SEM is shown). Data were analyzed with one-way ANOVA with Durnett correction, **p<0,01, ***p<0,001. **D.** Phase-contrast microscopy of colon and pancreatic cancer organoids derived from tumor patient's tissues. Representative pictures of the organoid culture morphology post treatment with MKT-077, CB6644, Selinexor and combinations of inhibitors for viability test shown in Figure 3 E-F. Scale bar 400µm.

### Supplementary Figure 4

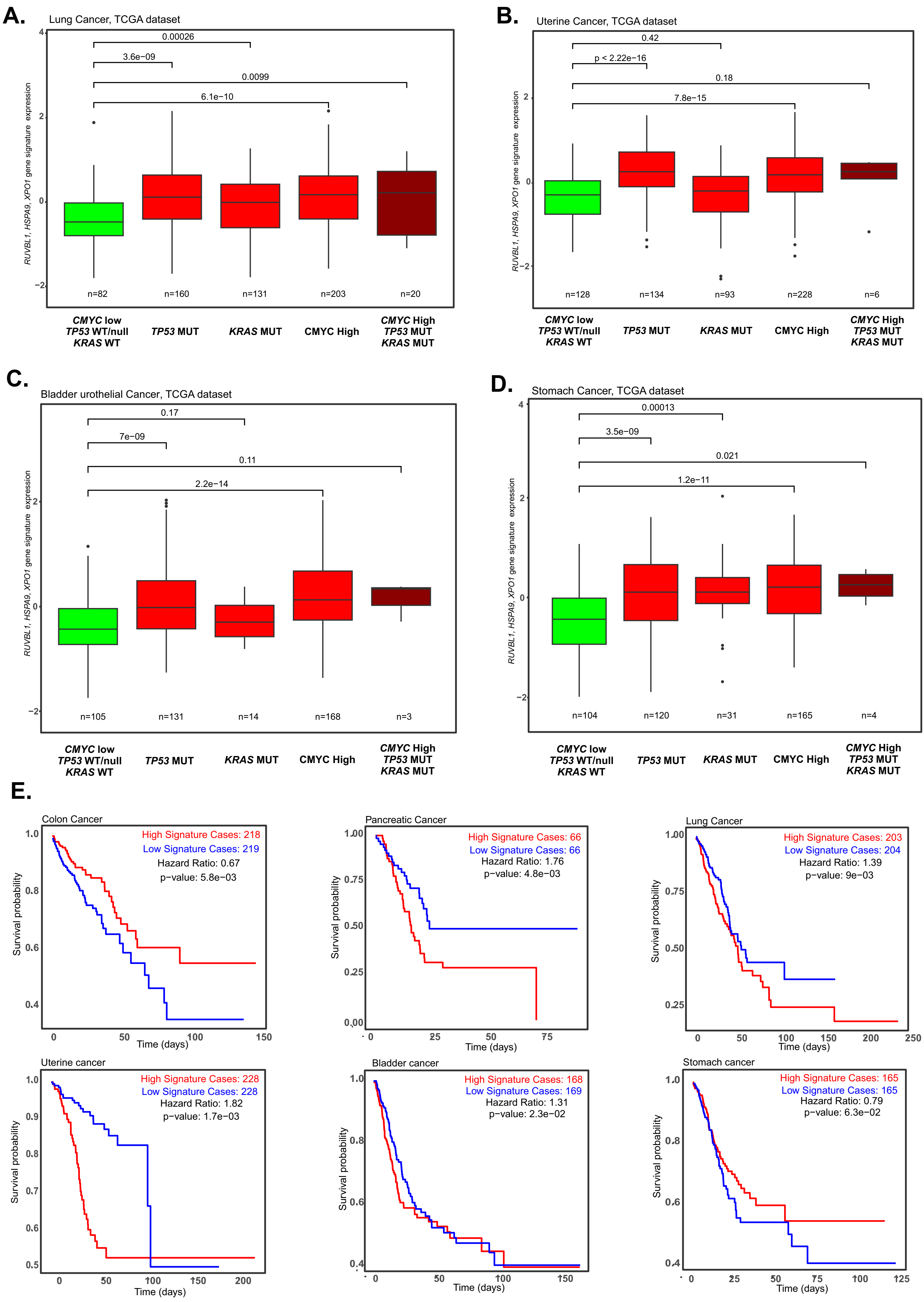

#### F. Figure 4 colon cancer samples:

| Patient: | Tumor tissue: | Diagnosis: | Oncogenes: |
|---|---|---|---|
| CC 1 | 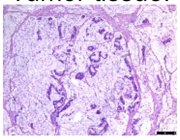 | Mucinous<br>cecal<br>adenocarcinoma<br>pT4a, N2a,<br>M1 | TP53 R273H<br>KRAS WT<br>CMYC low |
| CC 2 | 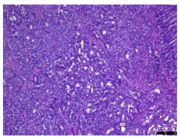 | Colon<br>adenocarcinoma<br>G2<br>pT4a, N2b<br>M1a | TP53 WT<br>KRAS WT<br>CMYC high |
| CC 3 | 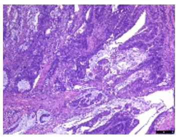 | Colon<br>adenocarcinoma<br>G2<br>pT2, N1b<br>R0 | TP53 WT<br>KRAS WT<br>CMYC high |
| CC 4 | 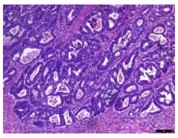 | Colon<br>adenocarcinoma<br>G2<br>pT3, N0<br>R0 | TP53 WT<br>KRAS WT<br>CMYC low |
| CC 5 | 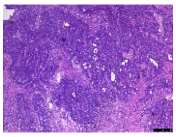 | Adenocarcinoma of the<br>rectosigmoid flexure<br>G2<br>pT2, N1a<br>R0 | TP53 212STOP<br>KRAS WT<br>CMYC low |
| CC 6 | 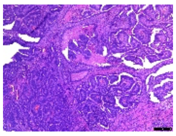 | Cecal<br>adenocarcinoma<br>G2<br>pT2, N1b<br>R0 | TP53 WT<br>KRAS WT<br>CMYC high |
| CC7 | 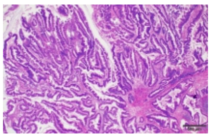 | Colon<br>adenocarcinoma<br>G1<br>pT3, N0<br>R0 | TP53 WT<br>KRAS WT<br>CMYC high |
| CC8 | 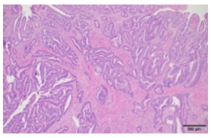 | Colon<br>adenocarcinoma<br>G1<br>pT2, N0<br>R0 | TP53 R282W<br>KRAS G12C<br>CMYC low |
| CC9 | 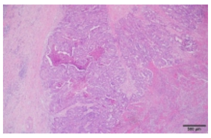 | Colon<br>adenocarcinoma<br>G2<br>pT3, N0<br>R0 | TP53 WT<br>KRAS WT<br>CMYC high |
| CC10 | 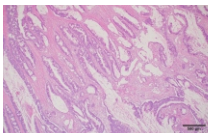 | Colon<br>adenocarcinoma<br>G2<br>pT3, N0<br>R0 | TP53 WT<br>KRAS WT<br>CMYC high |
| CC11 | 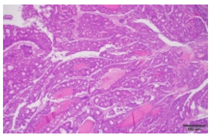 | Colon<br>adenocarcinoma<br>G2<br>pT3, N1b<br>R0 | TP53 WT<br>KRAS G12D<br>CMYC low |
| CC12 | 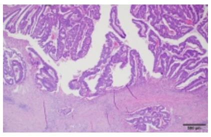 | Colon<br>adenocarcinoma<br>G2<br>pT3, N0<br>R0 | TP53 N239S<br>KRAS G12D<br>CMYC low |
| CC13 | 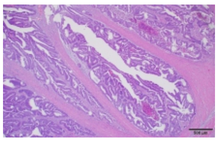 | Colon<br>adenocarcinoma<br>G2<br>pT3, N1a<br>R0 | TP53 WT<br>KRAS WT<br>CMYC high |
| CC14 | 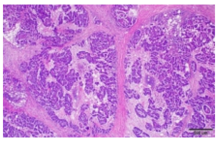 | Colon<br>adenocarcinoma<br>G3<br>pT4a, N2b<br>R0 | TP53 WT<br>KRAS WT<br>CMYC low |
| CC15 | 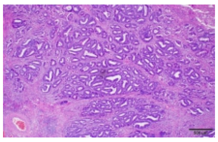 | Colon<br>adenocarcinoma<br>G2<br>pT3, N0<br>R0 | TP53 WT<br>KRAS WT<br>CMYC high |
| CC16 | 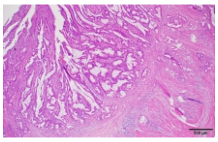 | Colon<br>adenocarcinoma<br>G2<br>pT3, N2a<br>R0 | TP53 R248Q<br>KRAS WT<br>CMYC low |
| CC17 | 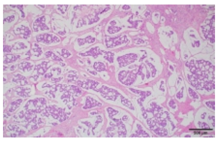 | Colon<br>adenocarcinoma<br>G2<br>pT3, N0<br>R0 | TP53 WT<br>KRAS WT<br>CMYC high |
| CC18 | 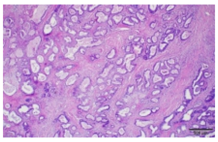 | Colon<br>adenocarcinoma<br>G2<br>pT3, N0<br>R0 | TP53 WT<br>KRAS WT<br>CMYC high |
| CC19 | 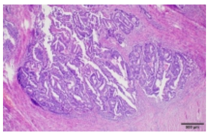 | Colon<br>adenocarcinoma<br>G2<br>pT2, N0<br>R0 | TP53 R258K<br>KRAS G12D<br>CMYC low |
| CC20 | 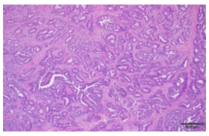 | Colon<br>adenocarcinoma<br>G2<br>pT2, N0<br>R0 | TP53 R273H<br>KRAS WT<br>CMYC high |
| CC21 | 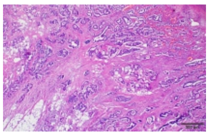 | Colon<br>adenocarcinoma<br>G2<br>pT3, N0<br>R0 | TP53 WT<br>KRAS WT<br>CMYC low |
| CC22 | 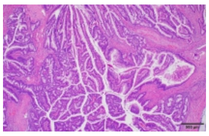 | Colon<br>adenocarcinoma<br>G2<br>pT3, N0<br>R0 | TP53 WT<br>KRAS G13D<br>CMYC low |
| CC23 | 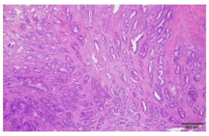 | Colon<br>adenocarcinoma<br>G2<br>pT3, N1b<br>M1 | TP53 R175H<br>KRAS WT<br>CMYC low |
| CC24 | 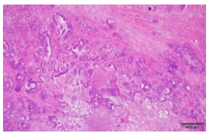 | Colon<br>adenocarcinoma<br>G2/G3<br>pT4a, N1b<br>M1 | TP53 WT<br>KRAS WT<br>CMYC low |

#### G. Figure 3 organoid colon cancer samples:

| Patient: | Tumor tissue: | Diagnosis: | Oncogenes: |
|---|---|---|---|
| CC25 | 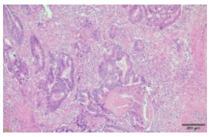 | Rectal<br>adenocarcinoma<br>G2<br>pT3, N0<br>R0 | TP53 H179Y<br>KRAS WT<br>CMYC high |
| CC26 | 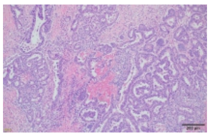 | Colon<br>adenocarcinoma<br>G2<br>pT4a, N0<br>R0 | TP53 C135F<br>KRAS G12D<br>CMYC high |
| CC27 | 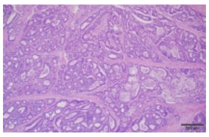 | Colon<br>adenocarcinoma<br>G2<br>pT3, N0<br>N0 | TP53 WT<br>KRAS V9F<br>CMYC high |
| CC28 | 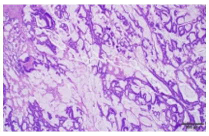 | Colon<br>adenocarcinoma<br>Gx<br>pT3, N2b<br>R0 | TP53 WT<br>KRAS G12D<br>CMYC low |

**H. Figure 4 pancreatic cancer samples:**

| Sample ID: | Tumor tissue: | Diagnosis: | Oncogenes: |
|---|---|---|---|
| PDA1 | 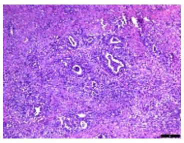 | Pancreatic adenocarcinoma<br>G3<br>pT2, N1<br>R0 | TP53 WT<br>KRAS WT<br>CMYC low |
| PDA2 | 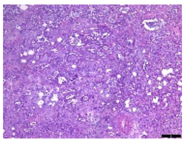 | Pancreatic adenocarcinoma<br>G2<br>pT2, N2<br>R1 | TP53 WT<br>KRAS WT<br>CMYC low |
| PDA3 | 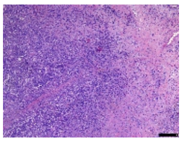 | Pancreatic adenocarcinoma<br>G3<br>pT2, N0<br>R0 | TP53 C238F<br>KRAS WT<br>CMYC high |
| PDA4 | 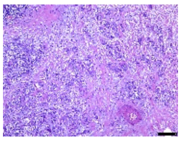 | Pancreatic adenocarcinoma<br>G3<br>pT3, N1<br>R1 | TP53 WT<br>KRAS G12R<br>CMYC low |
| PDA5 | 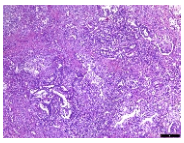 | Pancreatic adenocarcinoma<br>G2<br>pT3, N2<br>R1 | TP53 WT<br>KRAS G12D<br>CMYC low |
| PDA6 | 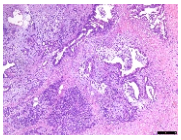 | Pancreatic adenocarcinoma<br>G2<br>pT3, N0<br>R1 | TP53 WT<br>KRAS WT<br>CMYC low |
| PDA7 | 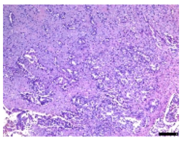 | Pancreatic adenocarcinoma<br>G2<br>pT2, N1<br>R1 | TP53 WT<br>KRAS WT<br>CMYC high |
| PDA8 | 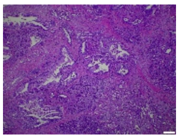 | Pancreatic adenocarcinoma<br>G2<br>pT2, N2<br>R1 | TP53 WT<br>KRAS G12D<br>CMYC low |
| PDA9 | 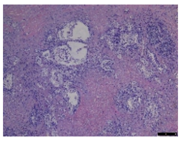 | Pancreatic adenocarcinoma<br>G2<br>pT3, N1<br>R1 | TP53 WT<br>KRAS WT<br>CMYC high |
| PDA10 | 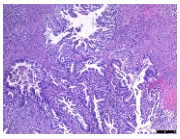 | Pancreatic adenocarcinoma<br>G2<br>pT3, N1<br>R1 | TP53 WT<br>KRAS WT<br>CMYC low |
| PDA11 | 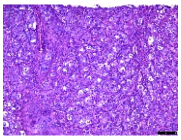 | Pancreatic adenocarcinoma<br><br>Liver metastsis | TP53 R193D<br>KRAS G12D<br>CMYC low |
| PDA12 | 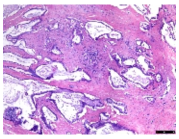 | Pancreatic adenocarcinoma<br>Gx<br>pT1c, N1, M1<br>R0 | TP53 WT<br>KRAS WT<br>CMYC low |
| PDA13 | 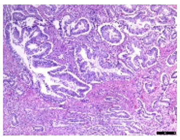 | Pancreatic adenocarcinoma<br>G1<br>pT1, N1<br>R1 | TP53 WT<br>KRAS WT<br>CMYC low |
| PDA14 | 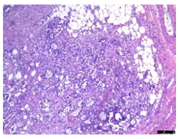 | Pancreatic adenocarcinoma<br>G1<br>pT2, N0<br>R1 | TP53 WT<br>KRAS WT<br>CMYC high |

**I. Figure 3 pancreatic cancer samples:**

| Sample ID: | Tumor tissue: | Diagnosis: | Oncogenes: |
|---|---|---|---|
| PDA15 | 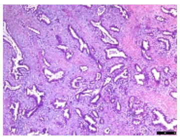 | Pancreatic adenocarcinoma<br>G2<br>pT3, N0<br>R1 | TP53 D281Y/H<br>KRAS WT<br>CMYC low |
| PDA16 | 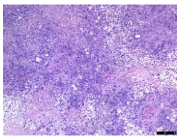 | Pancreatic adenocarcinoma<br>G3<br>pT2, N0<br>R0 | TP53 WT<br>KRAS G12D<br>CMYC high |
| PDA17 | 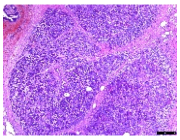 | Pancreatic adenocarcinoma<br>G3<br>pT3b, N1<br>R1 | TP53 H179N<br>KRAS WT<br>CMYC high |
| PDA18 | 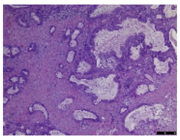 | Pancreatic adenocarcinoma<br>G2<br>pT2, N1<br>R1 | TP53 V193L<br>KRAS G12V<br>CMYC high |

**Supplementary Figure 4 A.** Comparative expression analysis of a 3-gene signature consisting of *RUVBL1, HSPA9* and *XPO1* in TCGA-derived patient samples of lung cancer (mean value of three genes in each patient was used to calculate the sample distribution in the box plot), stratified according to the listed *TP53, KRAS* (only point mutations) and *CMYC* expression status. The sample was included in "CMYC high" if the *CMYC* expression was above the *CMYC* average expression level for all the patients in the graph. T-test was used to obtain the indicated p-vaules. **B-D.** As in (A) for TCGA-derived patient samples of uterine, bladder urothelial and stomach cancers, respectively. **E.** Kaplan-Meier survival analysis of patients with the indicated cancer types, stratified according to the mean expression level of the *RUVBL1, HSPA9* and *XPO1* signature ("low" and "high"). Log rank test was used to obtain shown p-values for the difference between hifh and low expression levels of the signature. **F.** Hematoxylin and eosin staining histopathology photos, patient information/diagnosis and oncogene mutation/level status for *TP53*, *KRAS* and *CMYC* for colon cancer samples used in Figure 4A. **G.** As in (F) for colon cancer samples used to establish organoid cultures tested in Figure 3E. **H.** Hematoxylin and eosin staining histopathology photos, patient information/diagnosis and oncogene mutation/level status for *TP53*, *KRAS* and *CMYC* for pancreatic cancer samples used in Figure 4B. **I.** As in (F) for colon cancer samples used to establish organoid cultures tested in Figure 3F.

### Supplementary Figure 5

A.

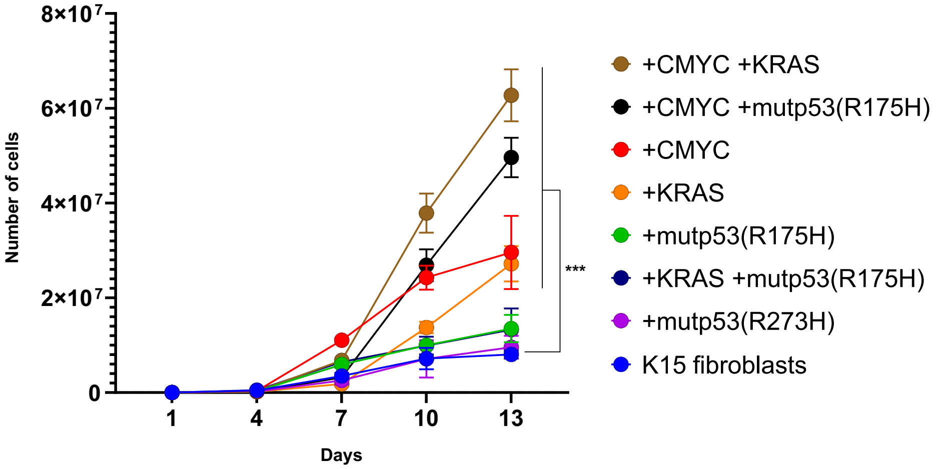


B.

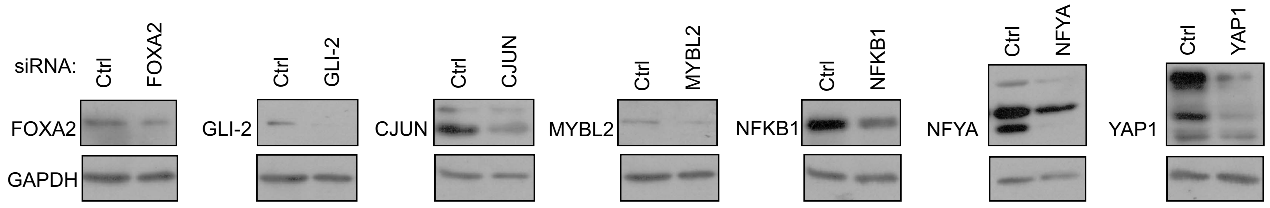


C.

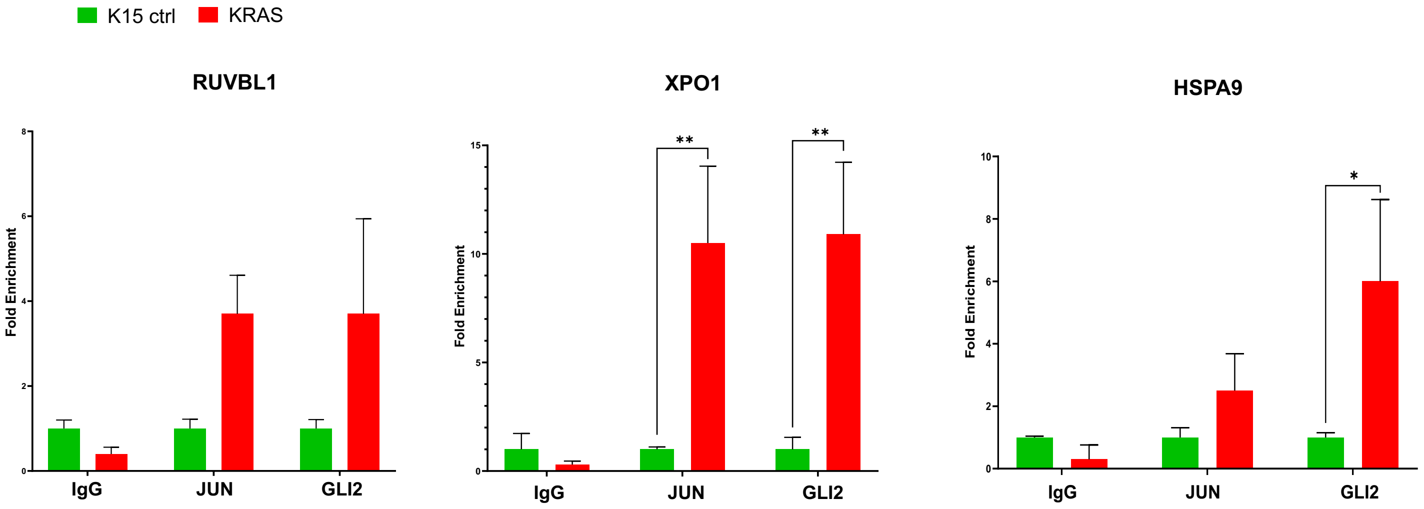

**Supplementary Figure 5 A.** Growth of K15 human fibroblasts with introduced listed single or double oncogenes. Cells were counted in 4-3 days intervals as shown in the graph. Means of n=2 are shown with SD, Two-way ANOVA test with Durnett correction, *** p<0.001. **B.** Western blots showing samples of K15 fibroblast cells with siRNA mediated silencing of indicated candidate co-factors of mutant KRAS or vs. non-speciofic silencing controls, setated to Figure 5G. Detected protein is indicated next to each blot with GAPDH detection as housekeeping control in all cases. **C.** Chromatin immunoprecipitaion-derived qPCR result of the promoter regions in the indicated genes, performed in either control K15 immortalized human fibroblasts or K15 fibroblasts with mutant KRAS stable overexpression. The PCR results were normalized to the levels of DNA precipitated in control K15 cells by each listed used antibody.

### Supplementary Figure 6

**A. Lung cancer cell lines:**

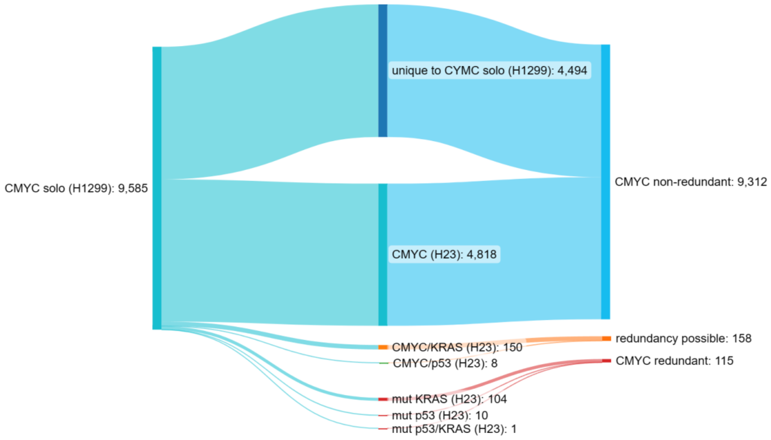


**B. Lung cancer cell lines:**

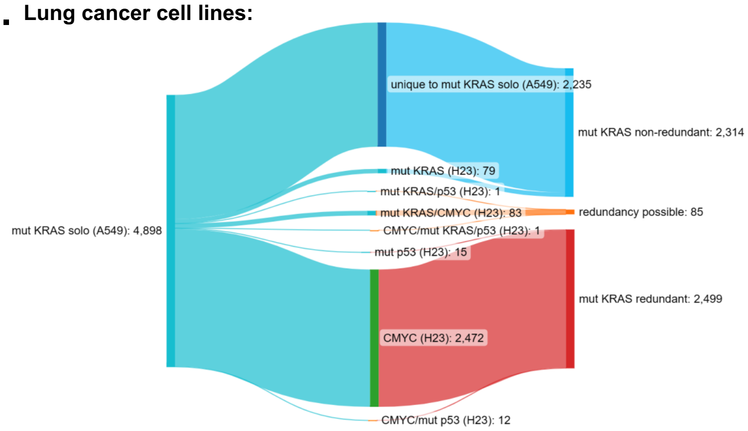


**C. Lung cancer cell lines:**

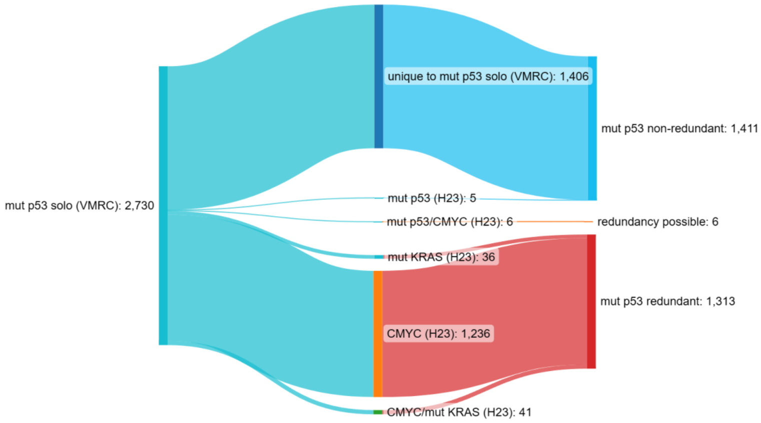


**D. Colon cancer patient datasets:**

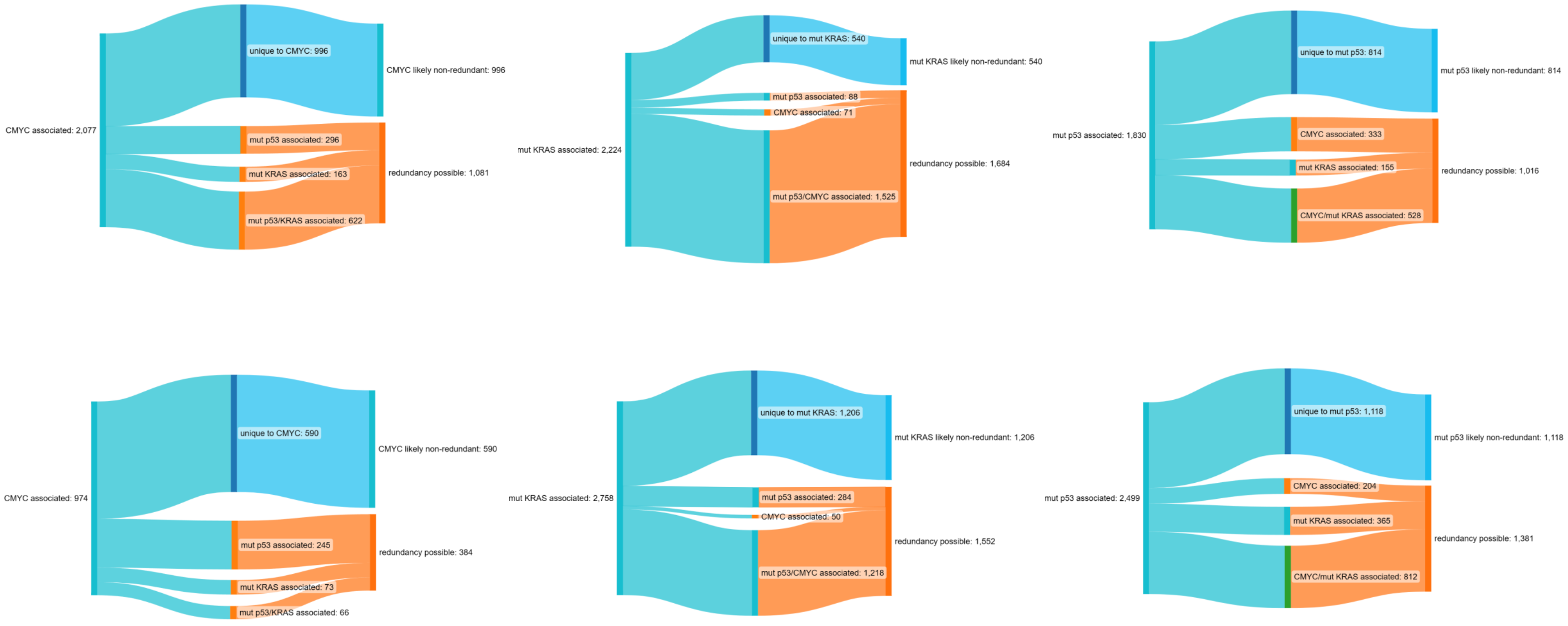

**Supplementary Figure 6 A-C.** Ribbon charts showing gene pools dependent significantly (FDR<0.05; only coding mRNAs) on mutant *TP53*, mutant *KRAS* or hyperactive *CMYC* (respectively) in lung cancer cell lines with single activated oncogenes (left end) shared with cell line with three co-activated oncogenes (middle), resulting in a specificity to the single oncogene (non-redundant genes), sharing with the co-expressed oncogenes (redundancy possible genes) or take-over by the co-expressed oncogenes from the program of the single oncogene (redundant genes; right end). The data on differentially expressed genes is derived from the CRISPR-Cas9 experiment in Fig. 1A and Supplementary table 2. **D.** Ribbon charts showing gene pools associated significantly (FDR<0.05; only coding mRNAs) to the presence of mutant *TP53*, mutant *KRAS* or hyperactive *CMYC* (left end in consecutive graphs) in TCGA-derived colon cancer patient dataset, shared or not with other co-activated oncogenes (middle), resulting in a specificity to the single oncogene (likely non-redundant genes), or sharing with the co-expressed oncogenes (redundancy possible genes). **E.** As in (D) for TCGA-derived lung cancer patient dataset. Data from D-E is summarized in Figure 6E.
